# The Cephalopod Reflectin Domain Specifically Chelates Cu(I), Inducing Photonic Assembly

**DOI:** 10.1101/2025.08.12.669908

**Authors:** Anjana R. Kammath, Irem Altan, Alison M. Sweeney

## Abstract

Reflectin proteins constitute the reflective layers of cephalopod skin responsible for camouflage and signaling. They are defined by repeated 25 a.a. motifs with conserved MDM tripeptides. The structure-function relations between these motifs and the proteins’ ability to assemble into a broad set of precisely controlled, optically resonant materials remain elusive. We observe that the MDM motif is also present in the Mets region of the Ctr copper transporter family, where it binds extracellular copper. We tested whether copper may also play an intrinsic role in reflectin protein assembly. Here we show that native reflectin-rich tissue has high concentrations of several transition metal ions, including Cu, and native protein will chelate additional Cu and Ni in the presence of EDTA and strong base. Experiments with two synthetic show reflectin motif peptides bind Cu(I) similar to Mets motif. Peptide-Cu(I) binding induces assembly with a stoichiometry dictated by Cu(I):peptide, resulting in a volume-spanning gel whose properties may inform our understanding of the emergence of optically resonant structures in the living system. We conclude that a core evolved function of reflectin domain is geometrically and thermodynamically specific assembly via Cu(I).

## Introduction

From a materials science perspective, structural color in living systems often comes from phase separation of a polymer melt secreted by cells into minimal surfaces (Fig. 1. In birds and insects, extracellular keratins, chitins, and waxes anneal into optically active gyroid and spinodal foam-type structures (e.g. ^2,3^). The physical principles of the emergence of minimal surfaces from polymer melts are well-understood (e.g., ^4^).

**Fig. 1:**
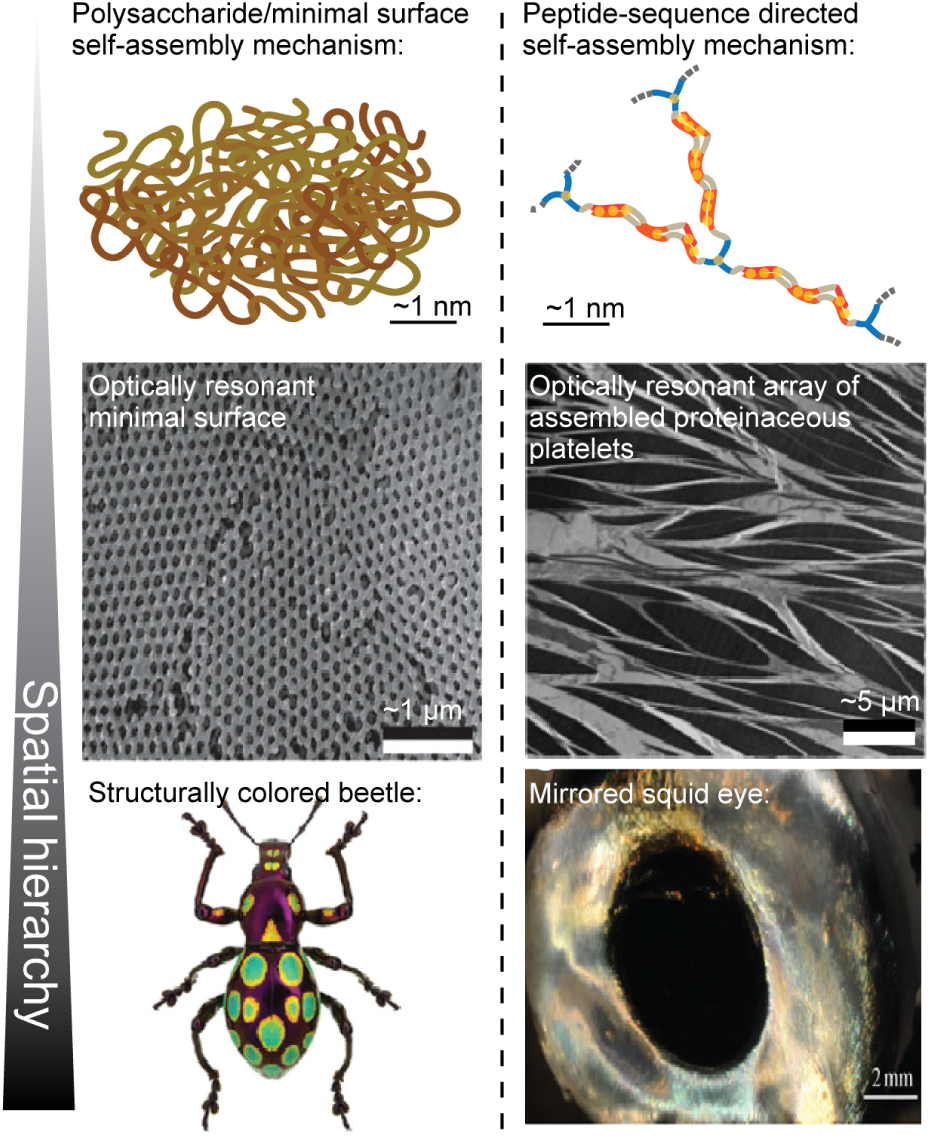
Material pathways to biophotonics. Left column: Many well-described biophotonic structures come from minimal surfaces that emerge from complex polymer melts. Right column: In contrast, cephalopod biophotonics emerge from platelet arrays composed of densely self-assembled reflectin protein. Beetle TEM and photo from. ^1^

However, the complexity of the polymeric mixtures employed by living systems and the sparsity of analytical tools for complex polysaccharides and glycoproteins makes it difficult, maybe impossible, to understand in chemical detail how these optically resonant structures emerge in living systems and are tuned by evolution. Similarly, the kinds of optical materials that may emerge through this route are limited to the optical geometries of minimal surfaces and phase separations.

In contrast, cephalopod skin is characterized by a layer of nano-scale, optically resonant materials in a wide array of geometries that differ markedly in geometry from minimal surfaces (Figs. 1, 2). Accordingly, the optical structures in cephalopods seem to have evolved a wider gamut of functions: guiding light, camouflage, signalling to conspecifics, and beam-shaping of bioluminescent emissions.^5–10^ The underlying arrayed structures are all composed of structural protein called “reflectin” that, in the living system, self-assembles into layered arrays of high-index protein-rich platelets and low-index intracellular fluid.

**Fig. 2:**
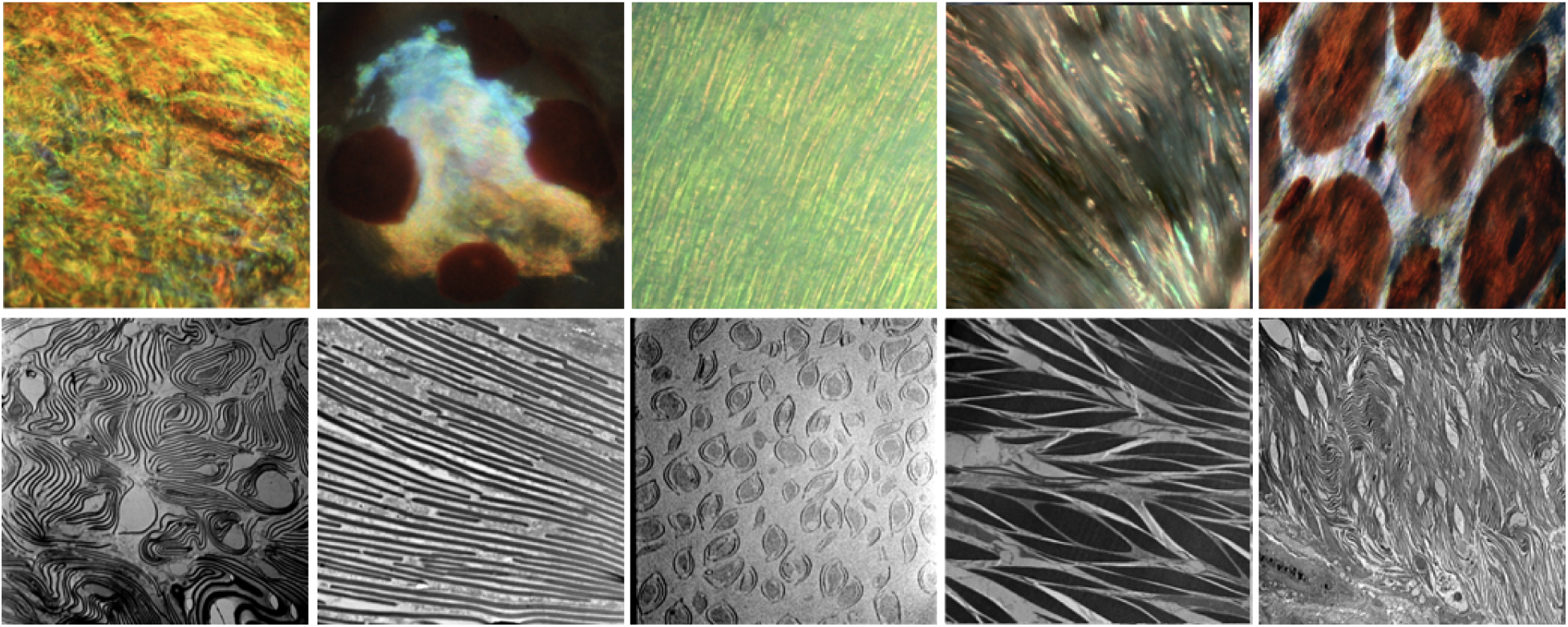
Examples of photonic cephalopod skin materials generated by reflectin proteins. Top row: reflected-light micrographs of squid skin, from left to right: *Chiroteuthis* eye reflector; *Chiroteuthis* photophore; *Galiteuthis* eye light guides; *Doryteuthis* eye reflector (tissue in this study); *Dosidicus* dorsal skin. Bottom row: Transmission electron micrographs in the same left-to-right order as the top row.

Because the cephalopod system assembles from a single structural protein motif with high evolutionary conservation,^11–14^ it presents a living photonic system with not only high functional flexibility, but also the prospect of understanding engineerable structure-function relationships of self-assembling photonic materials at a near-atomistic level of detail.

At the level of sequence, reflectin proteins seem to be intrinsically disordered.^15,16^ They have an unusual amino acid composition: rich in tyrosine, methionine, arginine, and tryptophan with a near-total absence of the hydrophobic residues alanine, isoleucine, leucine, and valine that are typically responsible for the hydrophobic collapse of canonical protein folding.^11–13^ Considered as polymers, reflectins are unremarkable weak polyelectrolytes.^16^ Considered as proteins, reflectins sit just near an order-disorder phase boundary, making an order vs. disorder classification of their final structure challenging.^16^ Across cephalopods, reflectins are characterized by two highly conserved motifs, an internal motif defined here as PER(Y/W/H)(M/F)DMS(N/G)(Y/W)(Q/C)MDM(Q/C/H)GR(W/Y)MD(M/R/S) (Q/W/Y)GR(Y/H/Q), and a similar but distinct N-terminal motif, defined here as MEPMSRM(S/T)MDF(Q/H)GRY(M/I)DS(M/Q)(G/D)R(M/I)VDP. These definitions are similar to those published previously but updated based on our own analysis of the most recent public genomic data. ^17–19^

In sequence aligments, the most conserved residues in these motifs are the residues and position of the tripeptide MDM, with 3–4 of these clusters repeated across the motifs.^11,16^ Though the N-terminal and internal motifs are clearly separately evolving entities, is difficult to say by eye what makes them different. The N-terminal motif has two highly conserved proline residues, one near each end; while the internal motif has a near-completely conserved ‘PER’ motif starting at the fifth position in the sequence. The sequence conservation of both motifs within the larger protein sequence strongly suggests that each has a distinct structural role in the larger reflectin assembly process. However, any functional role for these strongly conserved methionines is unknown.

Full-length reflectin proteins have a periodic sequence that begins with the N-terminal motif, followed by alternating aromatic-rich “linker” sequences and the internal reflectin motifs; all three of these sequence elements are roughly the same length. ^16^ In general, full-length reflectin proteins consist of an N-terminal motif and 3-4 repetitions of an internal motif, each coupled to an aromatic-rich linker region.

Previous *in vitro* work with recombinant reflectin has demonstrated some evidence of nano-scale self-assembly of small numbers of particles.^20^**^?^** ^-22^ These assemblies are reversible, suggesting that a route exists to dynamic assembly and disassembly of the higher-order optical properties.^20,23,24^ However, we have little or no understanding of the structure-function relationships linking the conserved secondary structure of the reflectin domain to the tertiary and quaternary structures implicated in assembly.

In considering the strong conservation of methionines in the N-terminal and internal reflectin motifs, we observed that it bears similarity to the Mets motif, which is part of the Ctr family of transmembrane copper transport proteins responsible for transporting copper ions from the extracellular milieu into the cytoplasm.^25–27^ We analyzed the reflectin motif in the context of all known Mets motif variants^26^ and found that reflectin clearly fits the Mets definition (Fig. S1).

Therefore, we investigated the hypothesis that, like the Mets motif, reflectin binds Cu and/or other transition metal ions. We discovered that reflectin proteins and synthetic peptides of reflectin motifs specifically bind Cu(I). Binding Cu(I) induces cross-linking and assembly of synthetic peptides of both the N-terminal and internal motifs described above. We also find significant concentrations of transition metals in native reflectin tissue, and that native reflectin protein reduces Cu(II) to Cu(I). After this auto-reduction, there is subsequent binding of Cu(I) with a mole ratio of ∼9 Cu(I) ions per protein. Similarly, we find that the synthetic motifs are capable of binding Cu(I) ions but not Cu(II) ions, as is also true of the Mets motif.^27^

When synthetic peptides of the internal and N-terminal reflectin motifs bind Cu(I), both peptides undergo structural shifts. However, the two motifs subsequently undergo markedly different assembly processes. The internal motif exhibits fractional binding stoichiometry, followed by a continuous assembly process that ends in a transparent gel. In contrast, the N-terminal motif shows integer stoichiometry of 3 peptides per 1 Cu(I) ion. We conclude that a core function of the conserved reflectin motif in the native system is to coordinate Cu(I). We infer that this metal coordination likely plays an important role in the further hierarchical assembly of these proteins into nanoscale structures capable of useful, evolved photonic resonance.

## Materials and Methods

### Peptide synthesis

We synthesized peptides representing the most evolutionarily conserved regions of the reflectin protein using a commercial service (GenScript, Piscataway, NJ). Most published reflectin sequences contain a methionine-rich sequence at the N-terminus ^23^ that is similar to but distinct from the similarly methionine-rich internally repeated, conserved peptide. Therefore, we used two different synthetic constructs to characterize the behavior of both the N-terminus and the internally repeated, conserved motifs of the proteins. Our syn-thetic N-terminus peptide had the sequence MEPMSRMSMDFQGRYMDSMGRMVDPK (26 amino acids, and designated “N-terminus”), while our internal motif had the sequence PERWMDMSNYSMDMQGRYMDRWGRY (25 amino acids, and designated “motif” or “internal motif”). Both peptides were synthesized to ≥ 95% sequence purity with N-terminal acetylation and C-terminal amidation. The motif peptide was insoluble in neutral aqueous buffer but fully dissolved in a solution of pH 3.5–4. The lysine at the C-terminal end of the synthetic N-terminus peptide is not part of the genetically encoded sequence; we added it to our construct when an initial 25-aa peptide representing the native sequence was predicted to be very insoluble^28^ at both neutral and acidic pH, similar to the Mets motif.^26,27,29^ Following the addition of lysine, the N-terminal peptide exhibited complete solubility in aqueous buffer conditions at pH 7 and below.

### Animal acquisition and use

Specimens of the squid *Doryteuthis pealeii* were obtained from the scientific collection resource at the Marine Biological Laboratory (MBL) or through assistance from the routine ecological survey work of the Connecticut Department of Energy and Environmental Protection (CT DEEP). Animals obtained from MBL were sacrificed according to that institution’s animal care and use protocols before shipment. Squid do not survive the survey trawling method used by CT DEEP, so were stored on ice immediately after collection and dissected within 24 hours. We separated fresh tissue from the eye lens, gills, and silver eye covering into cryovials. This tissue was either immediately used fresh in experiments, or stored at -80 °C. Unlike the other tissues used, gill tissue is exposed to ambient ocean water in the mantle cavity, so was rinsed in 1X PBS solution before freezing at -80 °C. When using either fresh tissues or removing tissues from frozen storage, we weighed the samples to a precision of ± 0.1 mg.

### Preparation of stock solutions

Ultrapure water was thoroughly degassed and traces of oxygen removed using the freezepump-thaw method. ^30^ The lyophilized motif and N-terminus peptides were combined with ultrapure water with the pH values described above to prepare the peptide stocks and then diluted as per experimental requirement. The motif peptide stock required a few rounds of vortexing to achieve complete dissolution. Concentrations of the resulting solutions were assayed via absorbance at 280 nm using a spectrophotometer (Nanodrop, Thermo Scientific) using the extinction coefficients *ɛ*_motif_ = 15220 M^−1^cm^−1^ and *ɛ*_N−term_ = 1280 M^−1^cm^−1^,^28^ and relative molar masses of M_r_,_motif_ = 3303.7 Da and M_r_,_N−term_ = 3155.7 Da. The ascorbic acid stock was prepared by dissolving 99% L-(+)-ascorbic acid (Fisher) in ultrapure water. Metal ion stock solutions were prepared by dissolving CuSO_4_· 5 H_2_O, CoCl_2_· 6 H_2_O, FeSO_4_· 7 H_2_O, NiSO_4_ · 4 H_2_O, ZnCl_2_, or AgNO_3_ in degassed ultrapure water (all reagents from Sigma except FeSO_4_ · 7 H_2_O, from Merck). Concentrated Cu(CH_3_CN)_4_]PF_6_ (Aldrich) stock solution was prepared in degassed acetonitrile to reduce the final concentration of acetonitrile in the peptide solutions.^26,27,31^ Ethylenediamine tetra-acetic acid solution (EDTA, Fisher) was prepared by dissolving in ultrapure water and then adjusting the pH to 8 using concentrated sodium hydroxide.

### Matrix Assisted Laser Desorption Ionization Mass Spectrometry (MALDI-MS) assay of peptide-metal binding

To detect possible stable interactions between our synthetic peptides and metal ions, we used MALDI-MS (AXIMA Confidence Linear Reflectron, Shimadzu Scientific Instruments). Importantly, this technique does not require a column separation; initial attempts at electronspray ionization MS, which does require initial column separation, were unsuccessful when peptides precipitated in the column. Peptide samples were prepared by diluting the stock with 5 mM ascorbic acid while maintaining the pH at 3.5. Measurements of MALDI-MS spectra of the peptides were calibrated and optimized by a series of preparations in different matrices (*α*-cyano-4-hydroxycinnamic acid, CHCA; 2,5-Dihydroxybenzoic acid, DHB; and a combination of CHCA and DHB; matrices were provided by the instrument manufacturer) and varying peptide concentrations in the presence of 5 mM ascorbic acid. Conditions of CHCA matrix with 30 µM peptide optimized signal:noise for both motif and N-terminus peptides. We then combined 1 µL of each peptide solution with 0.8 µL of CHCA matrix and pipetted the mixture onto the instrument’s 384-well plate (2.8 mm I.D.).

Metal ions were introduced to the system by pipetting two equivalents of the metals Cu, Fe, Ni, Co, Zn, and Ag to the matrix containing the peptide. To allow for adequate peptide-metal binding equilibration in experiments in which metal ions were present while also avoiding further peptide assembly or methionine oxidation, we first introduced peptide in aqueous solution to the matrix, then added metal ions in solution to this mixture, and then immediately dried the sample plate *in vacuo*. We found that this technique optimized peptide-metal ion interactions while simultaneously preventing oxidation of the many methionine residues in the peptide and the formation of larger peptide assemblies. In metalbinding competition experiments, ten equivalents of one competing metal ion in the form of CoCl_2_ · 6 H_2_O, FeSO_4_ · 7 H_2_O, NiSO_4_ · 4 H_2_O, ZnCl_2_, and AgNO_3_were added with two equivalents of Cu(I) ion. We also measured a concentration series of one to ten molar equivalents of CuSO_4_ to estimate the stoichiometry of peptide-Cu binding.

We collected mass spectra using the instrument’s N_2_ laser at 337 nm, 130–140 eV, and 3.0 Hz in linear mode in the range of 100 to 7000 m/z. The resulting spectrum from each sample is a cumulative of 121 profiles, with two laser shots accumulated at each profile. We calibrated these spectra using the standard mixture provided with the instrument according to the manufacturer’s instructions (TOF MIX, Shimadzu). Data post-processing was performed with the instrument-provided Launchpad software (Shimadzu Biotech, Kratos Analytical Ltd.).

### Ascorbic acid copper oxidation-state assay

To characterize the kinetics of peptide-Cu(I) binding, we conducted an ascorbic acid oxidation assay^32^ as implemented by Rubino and colleagues.^26,27^ In a control experiment, we monitored the aerobic oxidation of 100 µM ascorbic acid at pH 4.5 across a concentration gradient of 8–24 µM CuSO_4_ via optical absorbance from 245–265 nm every five seconds for five minutes in a 10 mm-pathlength quartz cuvette (NanoDrop, ThermoFisher). We then fit a relationship of ln A_255nm_vs. time vs. [CuSO_4_] to the resulting data to determine the correlation between the rates of ascorbic acid oxidation and [Cu(II)] (Fig. S5).

Then, to determine the influence of the reflectin peptides on Cu(I) oxidation, a series of peptide solutions with concentrations from 0-20 µM were combined with 100 µM ascorbic acid and 8 µM Cu(II). The fraction of free Cu as a function of the total concentration of peptide can then be determined by comparison to the control experiment.

### Dialysis-based metal dissociation assay

We dissolved 169 mg of the native silver eye tissue in 10 mL 0.1 M NaOH solution at pH 12. Three mL of this sample were transferred to 3.5 kD molecular-weight cut-off (MWCO) dialysis tubing (SnakeSkin, ThermoFisher). We reserved one mL of this parent sample in a microcentrifuge tube as a zero-time point reference. The dialysis tubing was transferred to 50 ml of dialysis buffer consisting of NaOH at pH 12 with 10 mM EDTA. The dialysis buffer was removed, stored, and then replaced with a fresh solution at time intervals of 30 minutes, 2 hours, and 21.5 hours. After 24 hours of dialysis, the solution of dissolved tissue was removed from the dialysis bag and also stored. As a control, a reference NaOH solution at pH 12 was placed in dialysis tubing and run in parallel to the above experiment under the same conditions. We lyophilized the stored samples for 24 hours, resulting in a small amount of hygroscopic material. We dried this material to completion on a hot plate at ≥300 °C for 24-48 hours until samples were completely dry. These samples were prepared for inductively coupled plasma mass-spectroscopy (ICP-MS) analysis by weighing them into clean Teflon vials, digesting using distilled nitric acid and hydrogen peroxide, and evaporating to dryness. When completely dry, the samples were brought back into solution in 5% nitric acid (Sigma) with ^115^In (1 ppb) as an internal standard. ICP-MS spectra for Na, Mg, Ca, V, Cr, Fe, Ni, Cu, Zn, Mo, Au, and Ti were collected at the Yale Analytical and Stable Isotope Center (YASIC; Element-XR, ThermoFisher). The spectrum of the reference sample was subtracted from the experimental samples for each time point collected.

### Bathocuproine disulfonate Cu(I) assay

To determine the valence and stoichiometry of reflectin-Cu binding in the native silver tissue, we employed the modified bathocuproine (BCS) assay. ^33^ 59.5 mg of flash-frozen reflectinbearing silver tissue was dissolved in 1 mL of 0.02% w/v NaOH solution with 10 mM of L-cysteine hydrochloride to serve as a reducing agent to convert all Cu species to the Cu(I) oxidation state. A second solution was prepared with 10 mM CuCl_2_, 10 mM L-cysteine hydrochloride, 6 mM EDTA, and 20 mM BCS. Increasing volumes of the Cu solution were combined with a fixed volume of the tissue solution, resulting in a Cu concentration gradient from 0.25-4.4 mM, and incubated for an hour. A control experiment was conducted in parallel by combining the same Cu(I) concentration gradient with the same NaOH-cysteine solution.

After this incubation, UV-VIS absorption was measured for all samples, with the absorbance at 483 nm used for the BCS assay and absorbance at 280 nm used to determine the protein concentration; in particular, it was important to account for L-cysteine absorbance at 280 nm in the control samples (NanoDrop, ThermoFisher). The absorption of the control was subtracted from the absorption of the sample at 483 nm and the Cu(I) concentration of the samples was determined from the standard curve generated by the dilution of the parent Cu(I) solution.

A second experiment with the BCS assay was conducted to identify the ratio of Cu(I): Cu(II) bound to the protein in the native tissue.^33^ Reflectin tissue was incubated in 2% SDS in aqueous buffer with 10 mM EDTA for 2 hours. The resulting liquid was subjected to chloroform-methanol extraction to remove all metals from the native tissue.^34^ The chloroform-methanol-SDS buffer mixture was centrifuged at 8000 g for 5 minutes and the resulting pellet was resuspended in the SDS buffer. The concentration of the resulting solution was determined using A280.

We then added, in series, 500 µM Cu(II), 10 mM EDTA, and 5 mM BCS. The reason for adding EDTA in this assay is the highly reactive nature of Cu^(II)^(BCS). We found that in this experiment, in the absence of EDTA, this species immediately reduces to more stable Cu^(I)^(BCS)_2_ leading to false positive results. The additional chelating agent was therefore required for replicability. By this logic, protein first interacts with Cu(II) in the absence of EDTA and BCS, and excess unreacted Cu(II) is then chelated with EDTA before reaction with BCS, preventing false positives. We then prepared another protein solution sample as described above but with the addition of 10 mM L-cysteine. Excess L-cysteine converts all Cu(II) to Cu(I).^35^ In this version of the assay, all Cu should be present in the form of Cu(I) and available for possible binding to the protein in the solution. We also prepared a control experiment with the same parameters as above but with no tissue added to the SDS solution.

A time series of absorption at 483 nm was then measured over 24 hours for both the test solutions and the control solution. We subtracted the control solution absorbance from both experimental solution absorbances and used these results to calculate the ratio of Cu(I): Cu(II) bound to the protein using the Beer’s law relation *A*_1_*/c*_1_ = *A*_2_*/c*_2_, assuming identical extinction coefficients and pathlengths for all samples.

### Circular dichroism (CD) and tryptophan fluorescence spectroscopy

We characterized the secondary structure of the synthetic peptides in the presence and absence of Cu(I) using CD spectroscopy (Chiroscan V100, Applied Photophysics). We measured the peptide in the presence and absence of Cu(I) introduced to the system in the form of [Cu(CH_3_CN)_4_]PF_6_. In these experiments, 350 µL of 5 µM of either motif or N-terminus peptide were measured with and without [Cu(CH_3_CN)_4_]PF_6_. The concentration of [Cu(CH_3_CN)_4_]PF_6_ ranged from 0–40 µM. The solutions were placed in a 1-mm pathlength quartz cuvette at 25degC. Triplicate measurements of each sample were made in 1-nm wavelength increments in the wavelength range of 190–260 nm. Background spectra of ultrapure water with and without acetonitrile were collected using the same parameters and subtracted from sample measurements accordingly. Mean residue molar ellipticities as a function of wavelength were calculated from the observed millidegree measurements.

We characterized the intrinsic tryptophan fluorescence of both motif and N-terminal synthetic peptides with and without Cu(I) ions in the sample. Aqueous solutions of 5 µM of peptide and 5 µM of peptide with 10 µM of [Cu(CH_3_CN)_4_]PF_6_ were placed in a 1-cm pathlength quartz cuvette, and fluorescence of the samples was measured with an excitation of 295 nm and emission recorded from 310–450 nm^36^ from a slitwidth of 5 nm and temperature of 25°C (Fluorolog 3 Time-Domain Fluorimeter, Horiba). We recorded the emission spectrum of ultrapure water in the same manner and used this for background subtraction from the sample measurements.

### Nuclear Magnetic Resonance for peptide-Cu(I) ligation chemistry

700 µM of motif peptide was freshly prepared in degassed D_2_O (Sigma Aldrich) in a N_2_ environment and sealed in 3 mM high-throughput NMR tubes. ^1^H NMR spectra of this sample were measured with a 600 MHz NMR spectrometer using conditions of 5 s relaxation delay and 1024 scans (Agilent DD2). A fresh stock of [Cu(CH_3_CN)_4_]PF_6_ was also prepared under nitrogen using degassed deuterated acetonitrile (CD_3_CN, Sigma Aldrich) and the resulting concentration of Cu(I) was verified using the BCS assay. Two equivalents of Cu(I) were added to the NMR tube containing motif also under N_2_ atmosphere, and ^1^H NMR spectra were collected using the same conditions. The same measurement was again repeated after the addition of three equivalents of Cu(I). Mnova software (Mestrelab Research) was used for all data analysis. The solvent peak of D_2_O and chemical shifts of trace ethanol impurities (at 1.17 ppm) ^37^ were used to reference and align the final spectra as shown in Fig. 4.

### Isothermal Calorimetry (ITC)

We performed a series of ITC experiments to understand the thermodynamics of peptide-Cu interactions^38^ and possible subsequent peptide assembly processes (Nano ITC Isothermal Titration Calorimeter, TA Instruments). All experiments were conducted at 25°C. Because the CuSO_4_ · 5 H_2_O and ascorbic acid in the solutions used for ITC are reactive and susceptible to oxidation,^31^ the exact concentration of Cu(I) in each Cu(I)-containing solution was calibrated before each experiment using the bathocuproinedisulfonic acid disodium salt (BCS) colorimetric assay described above. This assay exploits the absorbance of Cu^(I)^(BCS)_2_ at 485 nm with the extinction coefficient *ɛ*_485nm_ = 12400M^−1^cm^−1^.^39^ To control for the possibility of a direct interaction between the acetonitrile present in the system with either peptide or Cu(I), we also tested the competitive binding of acetonitrile to Cu(I) in the presence of the motif peptide using NMR. The ^1^H NMR spectrum of 200 µM [Cu(CH_3_CN)_4_]PF_6_ in D_2_O with 2 mM TCEP was compared to the ^1^H NMR spectrum of 100 µM motif with 200 µM [Cu(CH_3_CN)_4_]PF_6_ in D_2_O with 2 mM TCEP. This allowed the quantification of free acetonitrile with respect to the TCEP peak; the data were consistent with all acetonitrile in the system in an unbound state (Fig S2).

To find the enthalpy of Cu(I) binding to the motif peptide, we prepared a solution of 10 mM CuSO_4_ · 5 H_2_O in water and diluted this to a concentration of 1.5 mM using 5 mM ascorbic acid in water. We then prepared a 156 µM solution of the motif peptide in 5 mM ascorbic acid in water and degassed this solution under vacuum for 10 minutes. We loaded 50 µL of the CuSO_4_ · 5 H_2_O titrant into the instrument’s syringe, and 200 µL of the peptide solution into the instrument’s analyte cell. The analyte cell was stirred at 300 rpm for 30 minutes before the beginning of each titration. We then injected 2 µL of titrant into the analyte every five minutes with a total of 25 injections, with each injection monitored for 240 s. This experiment was also repeated using a different strategy to prevent oxidation, using 1.5 mM [Cu(CH_3_CN)_4_]PF_6_ in 2 mM tris(2-carboxyethyl)phosphine (TCEP) and 156 µM solution of the motif peptide.^40^ To study Cu(I) binding to the N-terminus peptide (135 µM), we repeated the experiment with these parameters but used [Cu(CH_3_CN)_4_]PF_6_ (1 mM) as a source of Cu(I) and 2 mM TCEP in place of ascorbic acid in both the titrant and analyte solutions. We controlled for the non-specific effects of these titrations, specifically the heats of dilution and other mechanical effects, in matching control experiments. These background data were subtracted from all raw data characterizing the peptides.^41^

To estimate the thermodynamic parameters of binding enthalpy (ΔH), equilibrium binding constant (K_a_) and binding stoichiometry (*n*), the resulting data were fit using the inbuilt models in NanoAnalyze (TA instruments). The N-terminus peptide titration data were best fit using the “independent” model in NanoAnalyze (TA instruments). This independent model describes the interaction of *n* ligands with a macromolecule that possesses either a single binding site or multiple equivalent binding sites. The motif peptide data were best fit using a two-site sequential model coupled to the independent model.

We further probed the system’s kinetics by conducting an experiment in which we performed a single injection of 3 mM Cu(I) into a concentration series of motif peptide from 0–1100 µM, resulting in a dataset of independent Cu(I):peptide from 0–100. The heat evolution after this single injection of Cu(I) was monitored for 10 minutes. We observed a transparent gel present in the sample cell after these experiments.

### Gel Density Analysis

To better understand the gel that resulted from excess Cu(I) and a longer equilibration time, the final motif-Cu samples were retrieved from the ITC cell, and we measured the density of the transparent gel and remaining supernatant. Briefly, the samples were incubated at 4 C for 24 hours and centrifuged for 5 mins at 8000 x g. The wet and dry weights of the supernatant (sparse phase) and gel (dense phase) were recorded before and after lyophilization respectively. The packing fractions were calculated using the total volume of water estimated from the weight difference, and the total volume of protein was calculated using the standard specific volume for proteins (0.708 mL g^−1^), and the known starting concentrations.

### Dynamic Light Scattering

We used dynamic light scattering (DLS) to characterize the hydrodynamic radii of the synthetic peptide solutions with and without Cu(I). These samples were prepared in the same manner as for CD experiments described above. 20 uL of each peptide solution with and without Cu(I) was pipetted into individual wells of a 384-well plate (Fisher Scientific), using care to avoid introducing air bubbles. The instrument was calibrated using 30 µM a bovine serum albumin standard (albumin standard, Thermo Scientific). Time correlations in light scattering were measured using a plate reader (DynaPro Plate Reader II, Wyatt). The instrument’s DYNAMICS software (Wyatt Technology) was used to calculate the cumulative of 10 acquisitions and the corresponding autocorrelation plots. The software’s built-in filter was used to remove noise from the acquisitions before taking the cumulative.

## Results

### MALDI-MS

In a sample of motif peptide without Cu(I) or other metal ions, we observed the most intense MALDI-MS peak at m/z=3304.7, consistent with the peptide as synthesized plus a proton (Fig. 3a). There are five methionines in the motif peptide and we also observe peaks consistent with their sequential oxidation plus an additional proton to the right of the peptide parent peak (Fig. 3a). A peak consistent with a triply-charged motif with three protons at m/z=1102.2 is the only other significant feature observed in this no-metal peptide spectrum. The internal peptides also showed evidence of fractional binding events (multiple peptides bound to one or more Cu(I)) and likely reaction intermediates (one Cu(I) bound to one peptide) (Fig. S11a,b).

**Fig. 3:**
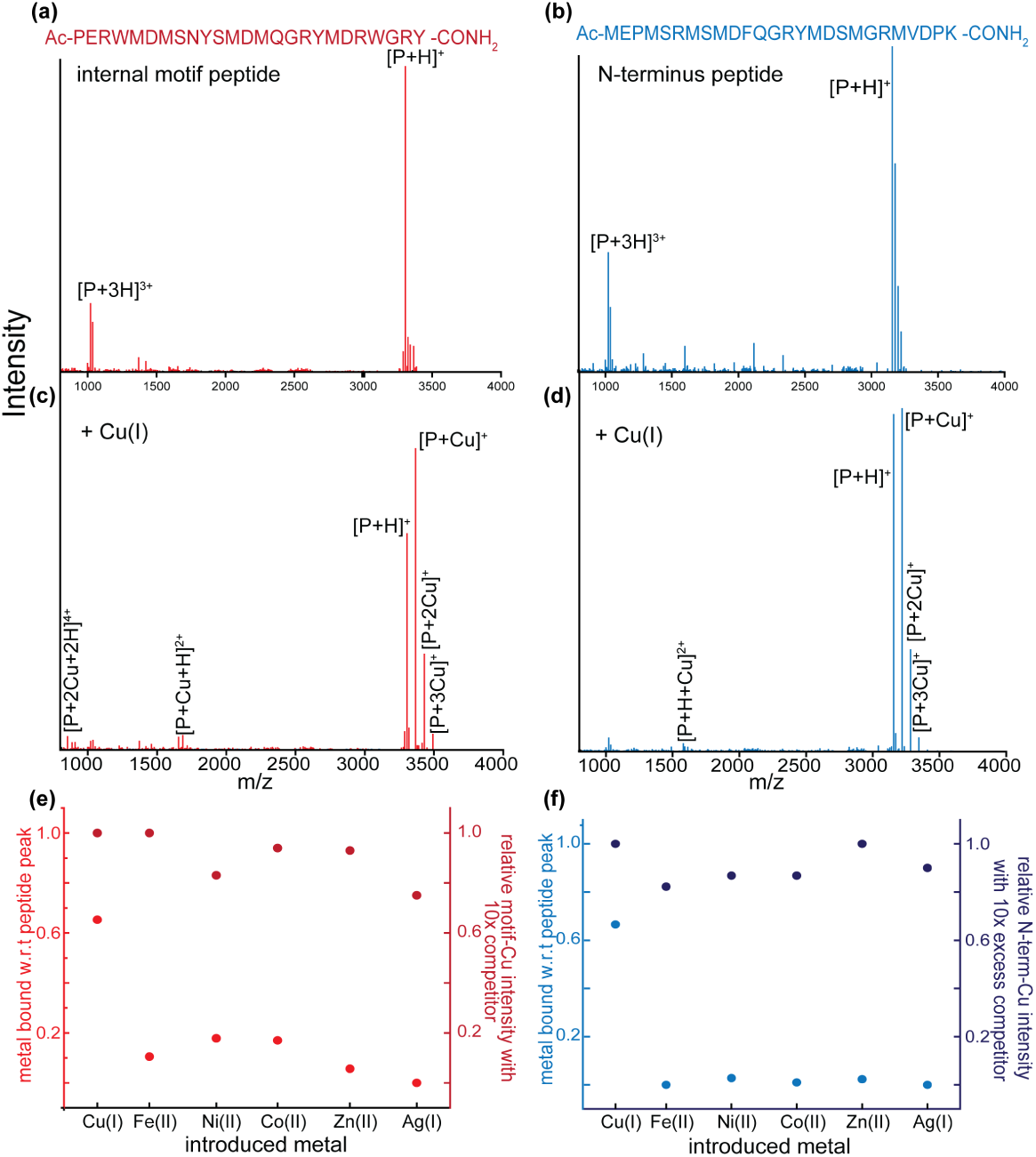
MALDI-MS spectra of motif and N-terminus peptides. The amino acid sequences of the peptides are shown at the top of each panel. **(a)** Motif peptide in the absence of Cu(I). **(b)** N-terminal peptide in the absence of Cu(I) **(c)** Motif peptide in the presence of Cu(I) **(d)** N-terminal peptide in the presence of Cu(I).

When N-terminus peptide was prepared without Cu(I), the most intense peak was at m/z=3156.7, and we also observed closely clustered peaks in this region, consistent with oxidation of the peptide’s methionines, as above (Fig. 3b). Unlike the internal motif, the N-terminal motif does not show evidence of fractional binding events in MALDI, but in context of ITC data may show reaction intermediates (Fig. S11c,d).

In a sample of motif peptide prepared with two equivalents of Cu(I) and 5 mM ascorbic acid, the peak corresponding to the parent motif ion was less intense than a new, right-shifted peak at m/z=3366.7 (Fig. 3c). This peak is consistent with the singly charged peptide bound to Cu(I). We also observe peaks at m/z=3429.5 and m/z=3492.5, which are consistent with the motif peptide with two and three Cu(I) adducts, respectively. These peaks are also absent in the no-metal measurement (Fig. 3c). These multiple-adduct peaks are also con-sistent with single-charge adducts formed by electron transfer without the loss of a proton.^42^ We also observe two minor peaks at m/z=1683.81 and m/z=857.8, which are consistent with a moiety with a charge of +2 resulting from the presence of a Cu and proton, and a moiety with a charge of +4, consistent with a charged motif bound to two Cu and two protons, respectively.

Similarly, when the N-terminal peptide was prepared with Cu(I), we observed a peak at m/z=3218.6 consistent with the peptide-Cu adduct with +1 charge (Fig. 3d). In this experiment, this peak was similar in intensity to the parent, unbound peptide peak at m/z=3156.7. We also observed peaks at m/z=3281.5 and z=3344.5 consistent with the N-terminal peptide bound to two Cu(I) and three Cu(I)(Fig. 3d). A less intense peak at m/z=1609.8 was also observed and is consistent with N-terminus peptide bound to a Cu(I) and proton(Fig. 3d).

We also observed that in the presence of Cu(I), the intensities of the peaks consistent with the oxidation of methionines markedly decreased in both peptide samples (Fig. 3c,d). The peaks in the motif-only sample consistent with methionine oxidation had intensities relative to the parent peak of 0.5, 0.12, 0.02, 0.05, and 0.02, for one to five oxidation occurrences, respectively.

However, in the presence of Cu(I), the relative intensities of the peaks representing methionine oxidation had lower relative intensities of 0.06, 0.08, 0.03, 0.05, and 0.1 respectively (Fig. S3a).

Samples containing the N-terminus peptide showed seven peaks consistent with the oxidation of each of the seven methionines in the sequence. The intensities of these peaks relative to the parent unoxidized peak ranged from 0.6 to 0.01 in the absence of Cu(I). When Cu(I) was present in the sample, only one of these peaks was present and was consistent with a single oxidation with an intensity relative to the parent peak of 0.2 (Fig. S3b).

To determine if this observed interaction between Cu(I) and the reflectin motifs was specific to this ion or a general ability of these peptides to bind metal ions, we conducted a metal-ion-binding competition study in which molar excess of a second metal ion was added to the sample along with Cu(I). In this assay, the peak associated with Cu(I)-bound motif peptide had an intensity 0.65 times that of the parent, unbound peptide peak (Fig. 3e). The peak intensities consistent with binding to the other metals assayed relative to the unbound parent peak were as follows: Fe(II), 0.1; Ni(II), 0.2; Co(II), 0.2; Zn(II), 0.07; Ag(I), below the detection limit.

When the N-terminal peptide was prepared with Cu(I) and competing ions, the Cu-bound peak had an intensity relative to the parent peak of 0.66. For the N-terminal peptide, all relative peak intensities consistent with the peptide binding the competing metal ion were less than 0.05 or undetectable (Fig. 3f).

In experiments where Cu(I) was introduced in a concentration series to both the motif and N-terminus peptides, we observed systematically increasing intensity of moieties consistent with 1:1, 1:2, and 1:3 peptide:Cu as a function of [Cu(I)], suggesting that copper binding increases with copper concentration (Fig. S4a,b). This trend was consistent across peptide:Cu conditions, however the overall intensity of the Cu-bound peaks decreased as the relative amount of copper increased from 1:1 to 1:3. In the context of our other results, we predict that this is due to the immediate gelation of the system in the presence of molar excess Cu(I) relative to peptide, and correspondingly reduced flight of the metal-bound moieties. In contrast, in samples in which ascorbic acid was absent, the presence of CuSO_4_ did not result in peaks consistent with any of the Cu-metal adducts described in Fig. 3 where ascorbic acid is present. Consistent with this, we also observed increased relative intensities of the peaks consistent with the oxidation of methionines in the motif and N-terminus peptides (Fig. S4c,d).

In total, these observations suggest that both the motif and N-terminus peptides coordinate Cu(I) via unoxidized methionine; the interaction of Cu(I) with both the internal motif and the N-terminal peptides suppresses methionine oxidation; and methionine oxidation inhibits Cu(I) binding. We also never observed a metal-bound peak with greater than 20% intensity of the Cu(I)-bound peak when peptides were exposed to both Cu(I) and a second metal ion in 10x excess, suggesting that while other metal binding events are possible, the most favorable and/or rapid binding event is with Cu(I). Further, the only peak in these experiments with greater than 0.5 relative intensity to the parent peak was the Cu(I)-bound peak, and the Cu(I)-bound peptide peak intensity is very similar to that of experiments with only Cu(I) present, suggesting that the presence of other metal ions does not significantly alter the affinity of the peptide for Cu(I). These results are consistent with a stable, specific coordination of the sulfur atoms of the peptide methionine residues with Cu(I) ions. ^43,44^

### Nuclear Magnetic Resonance Shows Peptide Methionine Thioethers Bind Cu(I)

The ^1^H NMR spectrum of the motif peptide showed a single chemical shift at 2.09 ppm, characteristic of the methyl group of methionine in the chemical context of random-coil proteins and peptides (Fig. 4c). ^45^ The integral of this region is consistent with the presence of six protons, and the region from 2.11–2.22 ppm accounts for the remaining methionine protons present in the peptide (Fig. 4). Upon the addition of two equivalents of Cu(I), we observed a downfield shift of the six-proton feature from 2.09 ppm to 2.10 ppm, and the appearance of four new peaks at 2.15, 2.17, 2.18, and 2.22 ppm. The peak at 2.10 ppm is still consistent with six protons, while the peaks at 2.15 and 2.22 ppm indicate three protons, and the peaks at 2.17 and 2.18 ppm, each indicate 1.5 protons. Therefore, at least one of the peptide’s methionine residues is in a highly asymmetric environment (Fig. 4b). We interpret these chemical shifts in the presence of Cu(I) to mean that in the presence of two equivalents of Cu(I), three out of five motif peptide methionine residues participate in direct Cu(I) coordination.

**Fig. 4:**
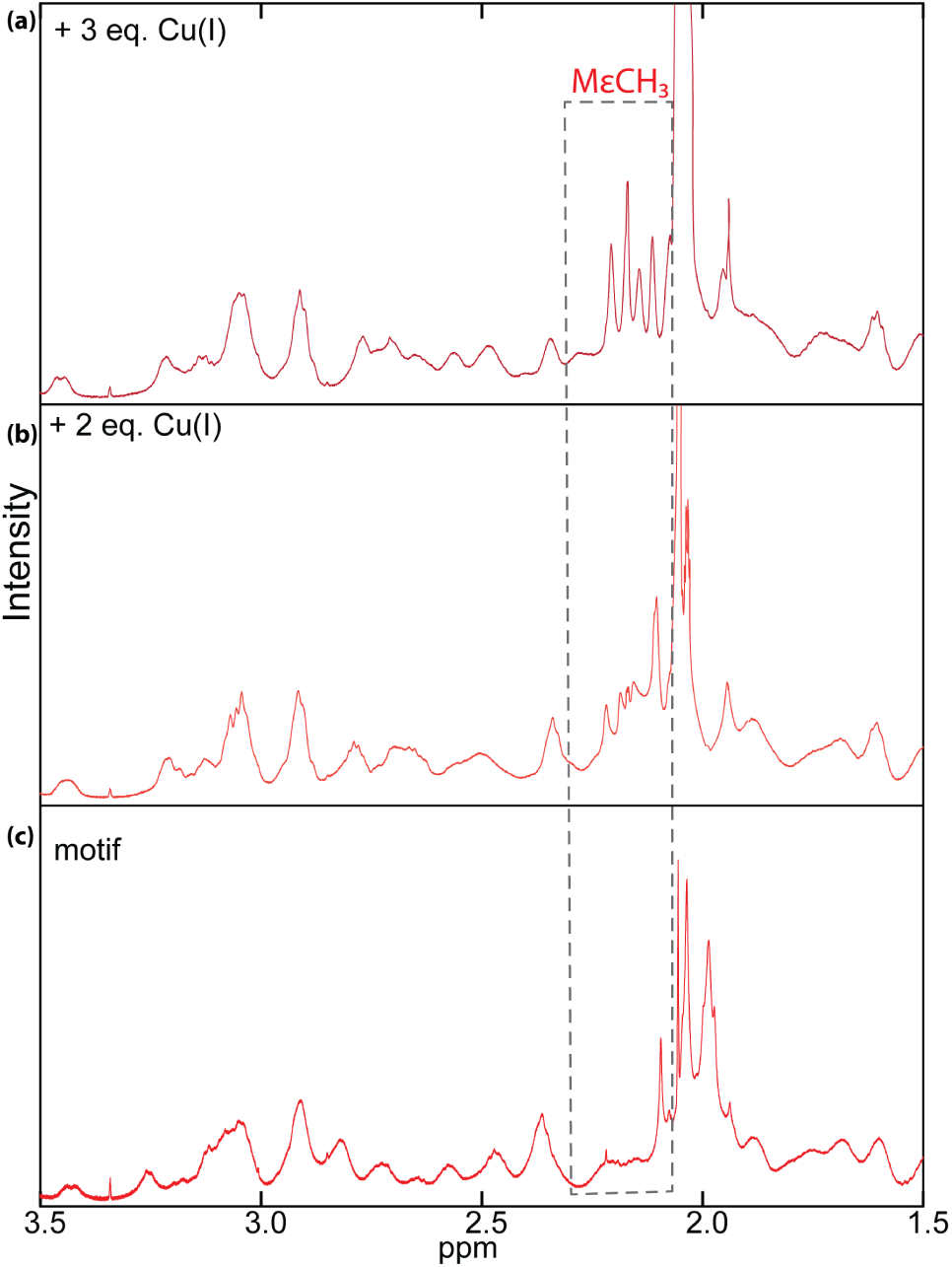
^1^**H NMR spectra of motif peptide in D_2_O. (a)** Internal motif peptide with three equivalents of [Cu(CH_3_CN)_4_]PF_6_ in CD_3_CN; **(b)** with two equivalents of [Cu(CH_3_CN)_4_]PF_6_ in CD_3_CN; **(c)** no added [Cu(CH_3_CN)_4_]PF_6_. The peaks associated with the methyl group of methionine shift downfield and undergo splitting consistent with the formation of methionine-thioether binding event(s).

Upon further addition of Cu(I) to three equivalents, the peak at 2.10 ppm shifts further downfield to 2.11 ppm, and new, well-defined peaks appear at 2.14, 2.17, 2.20, and 2.27 ppm (Fig. 4a). These peaks all have integrals consistent with three protons, showing that additional Cu(I) in the system causes additional shifts in the chemical environment of each of the motif peptide’s methionine. We also note that in the presence of two Cu(I) equivalents, the three methionines involved in Cu(I) binding shift upfield, while the other two methionines shift downfield (Fig. 4b). At two equivalents, the methionine peak at 2.27 ppm is also relatively broad, suggesting some amount of electron exchange between Cu(I) and methionine (Fig. 4b). This is consistent with the upfield shift of some of the methionine methyl groups being caused by aromatic ring currents shielding the methyl protons. The addition of further Cu(I) beyond three equivalents leads to the loss of peptide NMR signal, likely due to phase separation sedimentation of the peptide-Cu assembly described in more detail below.

### Ascorbic acid assay demonstrates complex reaction kinetics of peptide-Cu(I) interaction

The gradual addition of either peptide to the ascorbic acid-CuSO_4_ mixture results in a marked decrease in ascorbic acid oxidation, as monitored by absorbance at 255 nm (Fig. S5a). This is direct, specific evidence of Cu(I) binding the peptide.^46^ For both peptides, we observed an inflection in the decay rate after 200 s (Fig. S5a). When the N-terminal peptide was present, the rate of decay increased sharply at this time point, while when the motif peptide was present, the rate of decay increased gradually to the end of the assay. The kinetic rates generated from these exhibited a complex behavior and presented a challenge in fitting a simple binding relation to the data consistent with a constant average number of copper bound per peptide. Our data are more consistent with a mechanism in which there are fewer binding events per peptide as peptide concentration increases, possibly due to steric hindrance as peptide self-assembly commences at a critical peptide concentration (Fig. S5b,c). This result could also be due to an underestimation of free copper due to non-linear binding with concentration, or a combination of both effects.

### Dialysis and BCS Assay Show the Native Reflectin Tissue Contains Cu(I), Has High Affinity for Cu(I) and Can Auto-Reduce Cu(II) to Cu(I)

We dialyzed native tissue against a buffer of strong base and concentrated EDTA and measured the dialysate via ICP-MS over time to determine both the metal content and the qualitative affinity of the tissue for a range of transition metals (Fig. 5a,b).

**Fig. 5:**
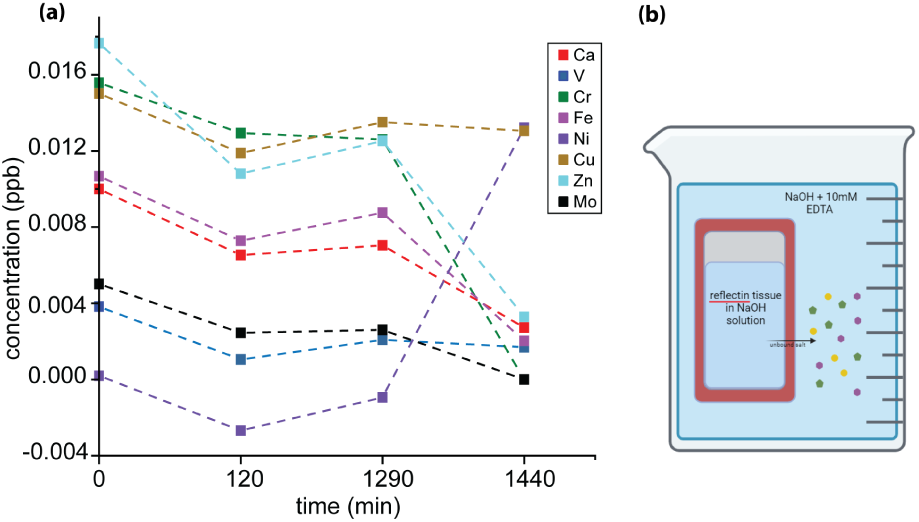
Metal ions in native silver tissue. **(a)** Dialysis-based assay of native-tissue metal ion affinity. Native silver tissue was placed in a dialysis cartridge and dialyzed against concentrated NaOH and EDTA. ICP-MS indicated the concentration of ions in the tissue over time. [Ca], [V], [Cr], [Fe], [Zn], and [Mo] were present in the native material and decreased severalfold over 24 hours. [Ni] and [Cu] increased in the native tissue after 24 hours. Inset: Schematic of the experimental design.

This buffer was designed to have a higher affinity for any given metal ion than a potential protein-ligand in the tissue. Over a 24-hour dialysis, the concentrations of calcium, vanadium, chromium, iron, zinc, and molybdenum in the suspension of native tissue decreased as expected from 0.010, 0.005, and 0.015 ppb, respectively, to approximately 0.003 ppb (Fig. 5a). [Na] remained unchanged at 0.0037 ppb, while [Mg] decreased from 0.06 ppb to 0.01 ppb (not shown). In contrast, [Ni] in the native tissue increased from undetectable to 0.015 ppb, suggesting that the tissue sample chelated the trace metal ion from the dialysis buffer (probably introduced to the system in trace amounts via the EDTA reagent or leached from glassware). Similarly, initial [Cu] in the tissue was 0.015 ppb, and did not change significantly (0.014 ppb) after 24 hours of dialysis. This result suggests that all the Cu present remained bound to the protein in the dialysis bag in spite of the presence of the highly metal-competitive dialysis buffer.

An ICP-MS analysis of digested native tissue revealed a range of metal concentrations elevated in reflectin tissue relative to lens tissue and gill tissue (Fig. S6a). In particular, V (80 µM) and Ni (2·10^4^ µM) were present in reflectin tissue at concentrations at least 10-fold higher than the control tissues. Cu ions were present at a concentration of 10^3^ µM in reflectin tissue, compared to a concentration of 3·10^4^ µM in the gill, where we expect to find high levels of copper, as they are an animal with a high respiration rate and a copper-based hemocyanin respiratory pigment.

We also performed a BCS assay on the native tissue, which confirmed the presence of significant concentrations of Cu(I) in the native reflectin tissue. The assay resulted in a linear increase in BCS absorption at 483 nm with respect to the Cu concentration. We fit the data to a line with a slope of 0.112 and y-intercept of 0.05 with *R*^2^ = 0.99. The *x*-intercept obtained from the fit was -0.44 mM for the protein sample and -0.0715 mM for the control (Fig. S6b). The difference in the x-intercept of the sample and control yielded [Cu]=368.44 µM in 42.54 µM reflectin protein. When extrapolated to the total protein in the tissue, this is a mole ratio of approximately 8.7 Cu ions per full-length reflectin protein. Given the concentration of protein in living tissue, this number of Cu(I) per protein is easily consistent with the value of 10^3^ µM in whole tissue obtained via ICP-MS.

We used a modification of the BCS assay to measure the oxidation state of Cu present in native tissue. The results from this experiment suggest that all the Cu present in the native tissue is in the +1 oxidation state and that the protein can reduce Cu(II) to Cu(I) and subsequently bind it, and that all Cu present in the system is bound to protein (Fig. S6c,d). In detail, we observed that in the presence of Cu(II) added to the system, BCS absorption monotonically increased for 2.5 hours, consistent with the reaction of the protein with Cu(II) and its conversion to Cu(I). Then, by repeating this experiment in the presence of excess cysteine, we were able to quantify the extent to which the initially observed reduction of Cu(II) to Cu(I) was due to native reflectin protein. Comparing a system with and without excess cysteine, after 2.5 hours the system with cysteine present had A483 of 0.934, while the solution with native protein and no additional cysteine showed an A483 of 0.11. Given that all 500 µM Cu(II) introduced to the system can be reduced by the excess cysteine, a comparison of the two results shows that the native protein sample contains 53.53 µM Cu(I). This is an approximate mole ratio of 8.9 Cu(I) per protein given an estimate of 27 kD average molecular weight in the complex native protein mixture, similar to the estimate of 8.7 Cu(I) ions per protein obtained independently above. After this initial 24-hour period, we observed that A483 decreased in both samples; in the protein-only solution it decreased from 0.11 to 0.064, and in the cysteine-containing solution it decreased from 0.934 to 0.641 while A280 remained unchanged in both. This longer-term change in the A483 signal in both samples suggests a second, slower mechanism of chelation of Cu(I) by the system, potentially via further protein assembly around slowly reduced Cu(I) ions.

### CD and tryptophan fluorescence spectroscopy show peptide structural rearrangements in the presence of Cu(I)

CD and tryptophan fluorescence analyses indicate large differences in the structure of the motif and N-terminus between the Cu(I)-bound and unbound states. In the Cu-free state, both synthetic peptides have CD spectra consistent with a random coil structure, exhibiting a strong minimum at 199 nm and 197 nm respectively (Fig. 6a,b). When [Cu(CH_3_CN)_4_]PF_6_ was added to a solution of the motif peptide, we observed a strong intensification of the minimum and blue shift from 199 nm to 197 nm. A shoulder also appeared at 220 nm (Fig. 6a). A prediction by BeStSel^47^ predicts this feature is due to the presence of helical structure. The structure of the N-terminal peptide behaved very differently in the presence of Cu(I). With this peptide, the addition of Cu(I) caused a strong positive increase in the initial minimum at 197 nm, and the appearance of a local maximum at 212 nm (Fig. 6b). The same structural prediction method suggests the appearance of *β*-turn structure in this peptide with Cu(I). With further titration of Cu(I) into the N-terminal peptide, this *β*-turn-containing conformation shifts to a weakly helical conformation given the two minima at 200 nm and 220 nm. In the presence of Cu(I), the addition of a high concentration of NaCl does not change the observed conformations (Fig. S7), suggesting that the Cu(I)-induced configurations of both peptides are non-trivially stable and not due to salting effects. In contrast, when Cu(II) was added to the system, we observed amyloid *β*-sheet structures in both peptides (Fig. S7).

**Fig. 6:**
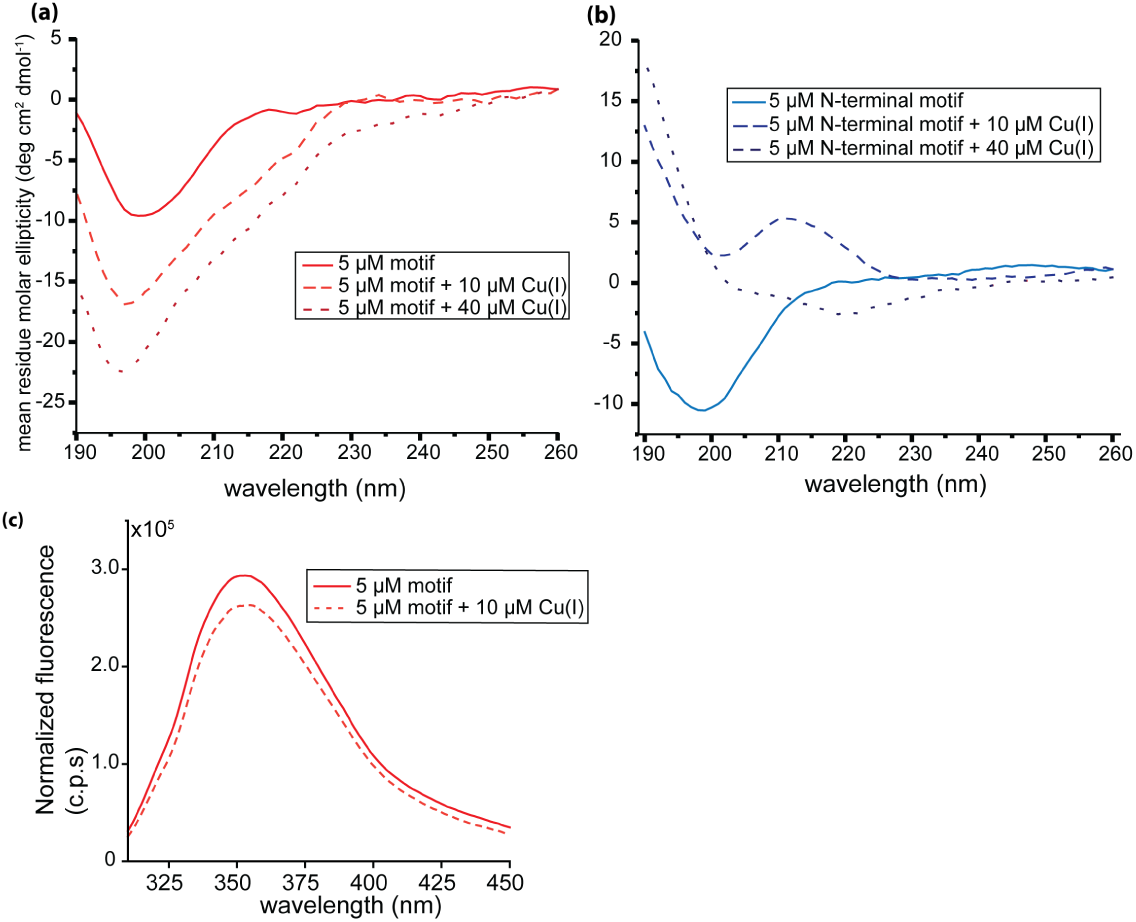
CD and tryptophan fluorescence show the peptide undergoes structural rearrangement on binding Cu(I). **(a)** CD spectrum of 5 µM motif peptide in the absence and presence of [Cu(CH_3_CN)_4_]PF_6_; **(b)** CD spectrum of 5 µM N-terminus peptide in water in the absence and presence of [Cu(CH_3_CN)_4_]PF_6_. **(c)** Tryptophan fluorescence spectrum of motif peptide in absence and presence of Cu(I).

The synthetic reflectin motif has two tryptophan residues in close proximity to two of the three regions of neighboring methionine sulfurs and the guanyl group of arginine that our data show are likely to mediate metal binding (WMDM at positions 4–7 and MDRW at positions 19–22). Tryptophan fluorescence of the synthetic motif peptide decreases in the presence of Cu(I), from a peak of 3 · 10^5^ counts per second in the Cu-free condition compared to 2.5 ·10^5^ counts per second in the presence of Cu(I) (Fig 6c). This result in concert with CD data suggests that sufficient structural rearrangement occurs upon metal binding to quench the tryptophans proximate to methionines. ^48,49^

### Isothermal calorimetry demonstrates that the N-terminus trimerizes and the motif peptide undergoes self-assembly in the presence of Cu(I)

We used isothermal calorimetry (ITC) to measure the thermodynamics of the peptide-Cu binding kinetics. The titration of Cu(I) into the motif peptide produced exothermic peaks on the order of -10^2^ µJ (−120 kJ mol^−1^) (Fig. 7a,b). These peaks increased in magnitude over the relative concentration range of 0 to 1 Cu:peptide from a value of -195.5 µJ (-65 kJ mol^−1^) to a value of - 355.3 µJ (-120 kJ mol^−1^). As the relative copper concentration further increased from one to three Cu:peptide, the magnitude of the peaks decreased from -355.3 µJ (-120 kJ mol^−1^) to -7.5 µJ (-3.5 kJ mol^−1^). As additional Cu(I) was added to the system, the thermogram then closely resembled the control, with the heats measured consistent with the heats of dilution in the control (Fig. 7a). The integration of this thermogram results in a sigmoid curve with an initial inflection point at 0.7 Cu:peptide. The sigmoid’s midpoint, indicating half-maximal activity, occurred at 1.2 Cu:peptide.

**Fig. 7:**
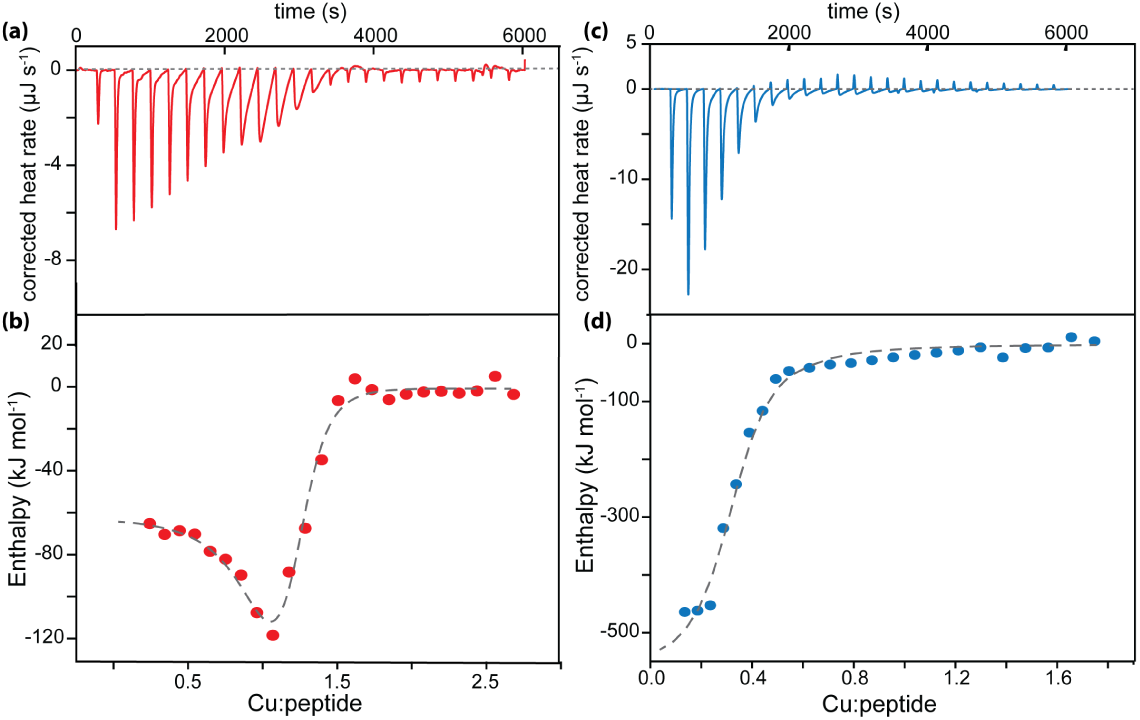
Isothermal Calorimetry (ITC) of Cu(I) and peptide binding and assembly. **(a)** Isothermal power as a function of the mole ratio of Cu(I):motif peptide. **(b)** Integrated heat as a function of Cu(I):motif peptide. The dashed line is the best fit generated by combining a two-site sequential model and an independent site model. **(c)** Isothermal power as a function of the mole ratio of Cu(I):N-terminal peptide. **(d)** Integrated heat as a function of Cu(I):motif peptide. The dashed line is the best fit generated from an independent site model.

Using standard fitting approaches incorporated in the ITC instrument’s software, this curve was best fit by merging a two-site sequential model with an independent binding model. The two-site sequential model fits the initial decrease in the sigmoidal curve, and the independent binding model fits the latter part.

When we used the N-terminus in this experiment, the titration of Cu(I) produced exothermic peaks with a magnitude of ∼ 10^2^ µJ, which diminished in size from -593 µJ (-449 kJ/mol) to -13 µJ (-8 kJ/mol) from 0 to 2 Cu:peptide (Fig. 7c,d). The remaining portion of the thermogram was consistent with the heat of dilution observed in controls (Fig. 7c). The integration of this thermogram produced a hyperbolic curve with an equilibration point at 0.3 Cu:protein. The data were well fit using an independent binding model in the instrument’s software, yielding the aforementioned equilibration point as binding stoichiometry (Table 1) and a binding enthalpy of ΔH = −570 kJ mol^−1^ (-136.23 kcal mol^−1^). The fit also gives K_A_ = 3.4 · 10^5^ M^−1^ and K_D_ = 2.9 µM.

**Table 1:**
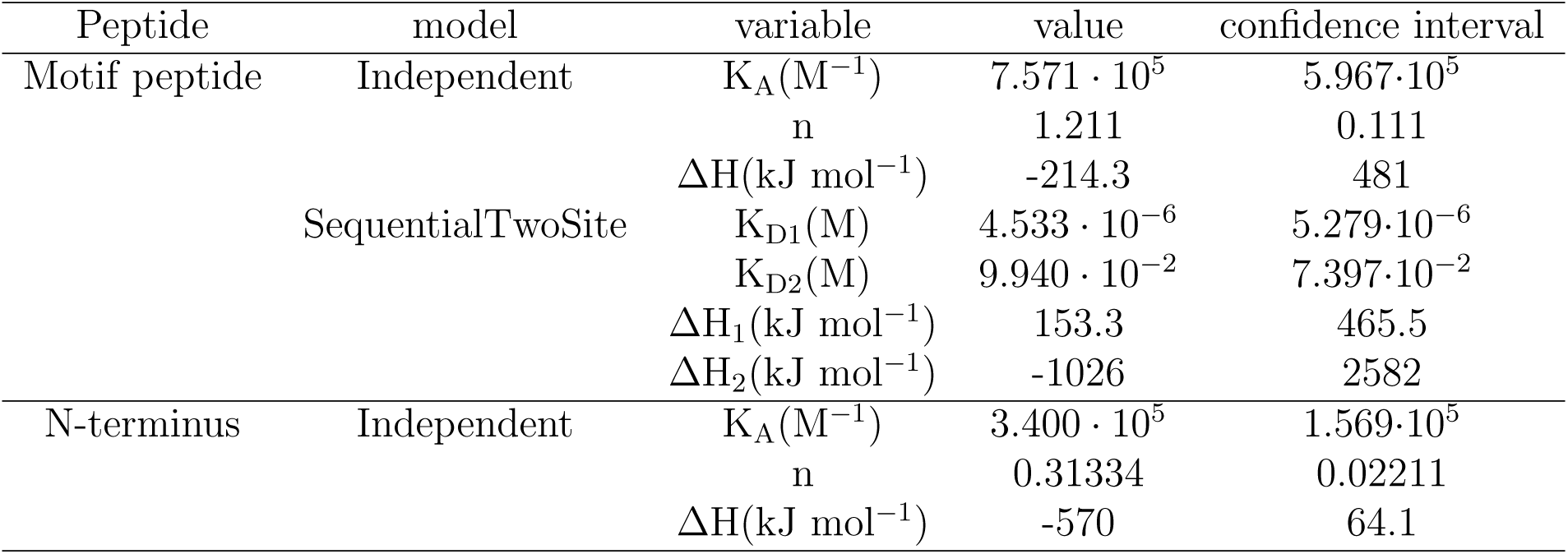
Peptide ITC fit data: The table summarizes the variables obtained from the fits described above for the titration of Cu(I) into the motif and N-terminus peptides. The rightmost column lists the values obtained for association constant (K_A_), stoichiometry (n), enthalpy (ΔH) and dissociation constant (K_D_). The confidence intervals are estimated for 95% confidence.

The thermodynamic parameters extrapolated from these fits are shown in Table 1. The Cu(I):internal peptide binding stoichiometry generated from the independent fit is 1.25 and the binding enthalpy is ΔH = −214.3 kJ mol^−1^ (−51.21 kcal mol^−1^).

The estimated association constant *K_A_* from this fit is 7.6 × 10^5^ *M* ^−1^ and the subsequent *K_D_* value calculated from the affinity is 1.3 µM. The titration of 1.5 mM Fe^2+^ and Cu^2+^ into the 50 µM motif solution served as controls for the experiments mentioned above (Fig. S5d,e). In these controls, each injection of the metal into the motif yielded a constant ∼ 0.2 µJ/s heat change, and an insignificant change in corresponding enthalpy (Fig. S5d,e).

We observed that both the magnitude and acceleration of heat evolution from the motif peptide system changed with increasing injection number and Cu:motif stoichiometry (Fig. S8a). In the initial Cu(I) injections, the acceleration of heat released by the system was high (12.1 kJ mol motif^−1^ s^−2^, rate parameter from fit - 0.045 s^−1^) with a similarly rapid return to baseline. In contrast, when the Cu(I):peptide ratio neared 1.7, the acceleration of the heat evolution was much less (2.38 kJ mol motif^−1^ s^−2^, 0.036 s^−1^) and returned to baseline similarly slowly (Fig. S8a). The total heat evolved in the early injections vs. the later injections decreased; early in the process, 262 kJ mol peptide^−1^ were released per injection, but 7 kJ mol peptide^−1^ at the end of the process. The initial rate of the reaction of Cu(I) with motif peptide is 4.5 ·10^−2^ s^−1^, and this rate decays to 1.6 ·10^−2^ s^−1^ when the reaction is near-saturated at a Cu:peptide value of 3 (Fig. S8a). These observations suggest that the initial behavior of the system is dominated by the rapid diffusion of Cu(I) ions and peptide-Cu binding events, while the later events require a slower process such as peptide rearrangements or peptide-peptide interactions in order for further peptide-Cu(I) interactions to occur. In contrast, the N-terminus peptide does not show similar rate complexity as the reaction with Cu(I) evolves (Fig. S8b). The initial rate of this reaction is similarly 4.5·10^−2^ s^−1^, but this reaction saturates after three injections and a Cu:peptide value of 0.3, after which we observe the rate of the reaction jump to ∼ 2 · 10^−1^*s*^−1^, consistent with simple heat of dissolution for Cu(I) (Fig. S8b, inset).

This observation of an initial fast event followed by a subsequent slower event in ITC motivated a subsequent ITC experiment to probe the system’s kinetics. Here we performed a single injection of a 25-fold molar excess of Cu(I) into a set of solutions of motif peptide of increasing concentration and monitored heat evolution over several minutes. In this experiment, we observed a sharp exothermic peak with a maximum at -160 µJ s^−1^ that required 150 seconds to relax to baseline (Fig. 8a).

**Fig. 8:**
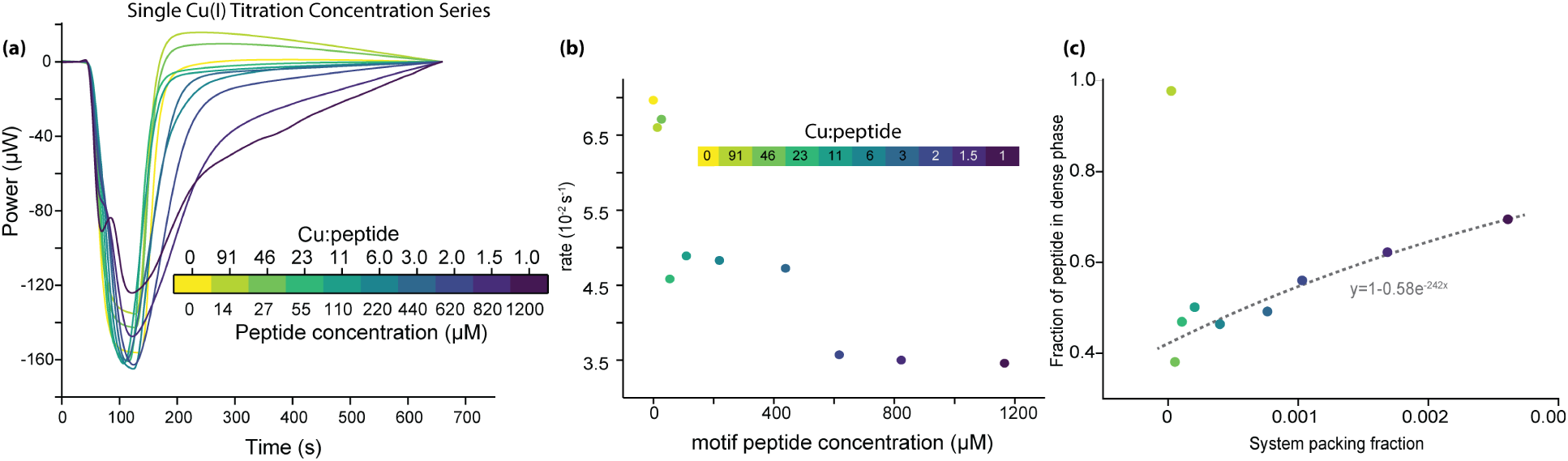
Single Cu(I) injection and long-time scale ITC. **(a)** ITC data for single injections of Cu(I) through a range of excess Cu(I):peptide molar ratios **(b)** Reaction rates calculated from data in **(a)**. **(c)** Phase separation of samples in **(a)**.

This phenomenon was particularly pronounced at low relative peptide concentrations (Fig. 8a). When Cu:peptide decreased to ≤1.7, we observed a change in the reaction kinetics. At these lower Cu:peptide ratios, we observed a shoulder at ∼60 s. This shoulder feature gradually lengthened in time with increasing relative peptide concentration. The exothermic heat release from each relative concentration measured fits well to a single exponential and an effective kinetic rate of the respective event was obtained (Figs. 8b). The kinetic rate reflects the gradual increase in the time evolution as it goes from 0.066*s*^−1^ for 14 µM motif peptide to 0.034*s*^−1^ for 1166 µM motif peptide. We interpret these shifts to indicate differences in the time required to shift from a copper-binding-dominated thermodynamic regime to a regime dominated by slower peptide-peptide interactions.

When we analyzed the phase separation we observed after this experiment (Fig. 8c), we found that the fraction of the peptide in the system in the gelled dense phase increased as a function of the total concentration of the system. In contrast, the same peptide remains fully soluble in a homogeneous transparent solution in the absence of copper at these same concentrations. The fraction of the peptide in the dense phase as a function of the total density of the system was well-fit by a simple relationship of *y* = 1 − 0.58*e*^−242^*^x^* where *y* is the fraction of the peptide in the dense phase and *x* is the packing fraction of the peptide. Extrapolating this relation to higher densities (that are not experimentally realizable due to the solubility of the metal-free peptide) predicts that the system undergoes complete phase separation to the dense phase at a packing fraction of ∼0.02.

### Dynamic Light Scattering (DLS) of peptide shows that the motif peptide undergoes sequential self-assembly in the presence of a Cu(I) concentration gradient

We titrated Cu(I) into solutions of the motif and N-terminal peptides, and in both cases, observed increases in the characteristic hydrodynamic radius with the addition of Cu(I) via DLS. In the absence of Cu, both the motif and N-terminus peptides display a time autocorrelation of 0.1–1 *µ*s indicative of particles *<* 1 nm in hydrodynamic radius, consistent with isolated peptide monomers (Fig. S9a,b). For the motif peptide, the time autocorrelation increased to 10–100 *µ*s as the Cu:peptide mole ratio increased from 0 to 4.8. When the mole ratio was greater than 0.7, the polydispersity of the motif-Cu solution increased, as indicated by the non-monotonic autocorrelation function. This result is consistent with a mixture of particles with hydrodynamic radii from 0.5–10 nm. Further addition of Cu further increases the polydispersity of the system, resulting in a system with some particles with hydrodynamic radii greater than 10 nm.

The N-terminus peptide showed a pronounced shift to longer autocorrelation times at a mole ratio of 0.5, with further additions of Cu causing further time delay in the autocorrelation function. This shift in autocorrelation accommodates the polydispersity in the particle size observed in the sample wells.

## Discussion

The myriad variety of optically resonant arrays found in cephalopod skin are all built from the structural protein reflectin. Though the protein class was discovered in 2004^11^, and is characterized by a highly conserved ∼25 aa motif with completely conserved methionine residues, to date we know very little about the structure-function relationships of this motif or the reason for its high degree of evolutionary conservation. We observe via several independent, complementary techniques that there are specific binding events between Cu(I) and synthetic peptides that characterize these evolutionarily conserved regions of reflectin proteins (Figs. 3, S5, 7), and that the native tissue is rich both Cu and other metal ions (Fig. S6). Similarly, CD analysis and tryptophan fluorescence show a large conformational change in both peptides upon Cu(I) binding, and the structures of the two peptides we analyzed differed markedly from each other both initially and upon binding Cu(I) (Fig. 6).

The binding event between Cu(I) and both peptides is specific and energetic; however, a surprising result of this work is that the thermodynamics of Cu(I) binding are entirely different between the two peptides. The internal motif peptide exhibits a sharp exothermic inflection at 1:1 Cu(I):peptide, followed by an endothermic release of heat that reaches half-saturation at around 3:2 Cu(I) peptide. This reaction then ceases to evolve heat as further Cu(I) is titrated into the system (Fig. 7b,d). In contrast, the reaction between N-terminal peptide and Cu(I) is always endothermic with a half-saturation around 1:3 Cu(I):peptide, and after this process is complete the reaction ceases to evolve heat.

Upon addition of Cu(I) to the internal motif peptide, we observed additional signals in the 2.1–2.3 ppm region corresponding to the *ε*-methyl group of methionine. This observation seems to confirm that the peptide’s methionines are primary participants in Cu(I) binding, likely via the methionine sulfur atom (Fig. 4). Further support of methionine-mediated binding comes from our observations that oxidation of methionines reduces Cu(I) binding in both the internal motif and N-terminal peptides (Fig. S3). We also observed that the BCS assay of this system is extremely sensitive to air exposure; the BCS assay was replicable only when extensive precautions were taken to degas all reagents and then performed in a nitrogen environment, thereby preventing methionine oxidation to the extent practicably possible.

The bonding energy of Cu + 2MeSMe → Cu(MeSMe)_2_ is estimated to be ΔH = −101.4 kcal mol^−1^ (−424.26 kJ mol^−1^).^50^ Our experimental finding in ITC for the motif peptide shows Δ*H* to be −214kJ mol^−1^ of peptide in an independent model, consistent with a single Cu-MeS bond per peptide. In the sequential two-site fit, we find an initial bond with Δ *H* = 153 *kJ mol*^−1^ of peptide and a second bond with Δ *H* = −1026 kJ mol^−1^ of peptide, consistent with two sequential Cu-MeS bonds per peptide (Table 1).

We infer from ITC that peptide-Cu(I) binding is most often an inter-peptide event given the fractional stoichiometry we observe for both motif and N-terminal peptides. Further, NMR shows that a thioether bond shifts upon introduction of Cu(I) to the system, and Cu(I) is often most stable in a tetrahedral geometry (Fig. 4).^51,52^

We also observe that the ‘MDM’ motif that is conserved in both peptides provides the correct bond angles and distances to create a site for inter-peptide Cu(I) center. In this binding geometry, the central aspartate residues are also necessarily oriented into solvent and away from the metal center in favorable fashion. Therefore, we speculate that in the metal-saturated system, Cu(I) may be coordinated by four methionines from two clusters of ‘MDM’ from separate peptides as illustrated in Fig. 9.

**Fig. 9:**
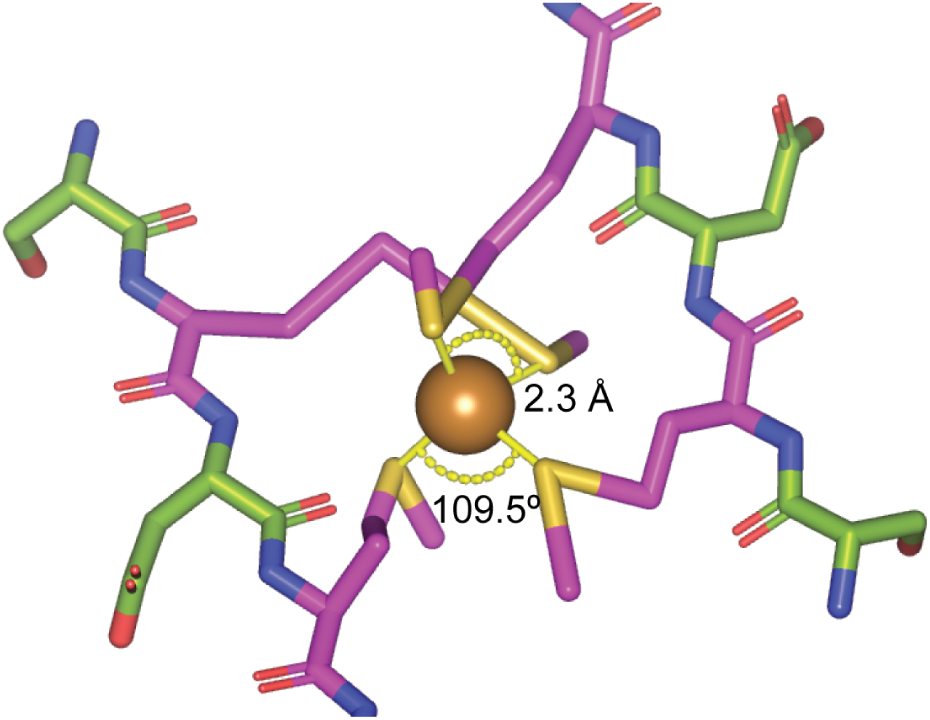
Data-informed hypothetical structure of Cu(I) bound in tetrahedral coordination to two MDM motifs in reflectin peptides. Methionine residues shown in magenta, sulfur atoms shown in yellow, Cu(I) shown in orange. Other residues shown in green. Distances and bond angles are labeled. This structure is consistent with the data presented here and has no steric clashes or unfavorable bond angles.

For insight into how this Cu(I) binding may inform the ability of these proteins self-assemble into optically resonant arrays *in vivo*, the differences in Cu(I) binding behavior between the two motifs are particularly interesting. Both show evidence of strong, specific binding with Cu(I) (Fig. 7) but not with other metal ions such as Fe(II) (Fig. S10). The internal motif exhibits an unusual fractional binding stoichiometry of 1.2 Cu(I):peptide that suggests that on average, two or three peptides are cross-linked by three Cu(I) ions. Knowing that methionine residues are primary participants in this binding, it is interesting to examine the pattern of methionines in the secondary sequence. The internal motif peptide has one MDM motif near the N-terminus, a second at the center, and an MDR motif near the C-terminus, providing three distinct potential sites within the peptide for potentiating Cu(I) coordination. If these three sites are capable of coordinating Cu(I) across separate peptides, this both accounts for the observed stoichiometry and suggests a mechanism for self-assembly into a network of occasionally branching strands. Our BCS characterization of native protein showed there are 8.9 Cu(I) per full-length protein in squid tissue (Fig. S6b). If each internal motif binds an average of 1.5 Cu(I) ions and a full-length protein has 3-5 internal motifs^11^**^?^**, our data suggest that proteins in the living tissue are saturated with Cu(I).

This hypothetical mechanism of self-assembly is qualitatively consistent with our observation that in the presence of excess Cu(I), the system of internal motif peptide alone will undergo phase separation into a sparse phase of peptide in solution and a dense phase that is a transparent gel (Fig 8c). Further, when injecting excess Cu(I) into a series of internal motif peptide solutions of increasing concentration, the evolution of heat in the system slows markedly, suggesting that as peptide concentration increases, the energy that dominates the system shifts from rapid Cu(I) binding events to slower peptide-peptide interaction and assembly events (Fig. 8a).

Surprisingly, though the N-terminal motif is equally methionine-rich and also exhibits assembly as observed in ITC, its assembly process appears to be completely disparate from that of the internal motif. Unlike the complex thermodynamics observed for the internal motif, the N-terminal motif exhibits a straightforward binding curve in ITC showing that three peptides are cross-linked by a single Cu(I) ion (Fig. 7c,d). Analysis of reaction rate as a function of Cu(I) concentration reveals an initial rapid, energetic event to the point of saturation (Fig. S8b), after which little further heat is evolved in the system. This is in marked contrast to the internal motif peptide which exhibits a continuous slowing of heat evolution with increasing [Cu(I)] which we were unable to saturate (Fig. S8a,b).

Unlike for the internal motif, inspection of the secondary structure of the N-terminal motif provides little insight into a possible mechanism for formation of stable trimers via binding Cu(I). Like for the internal motif, methionine oxidation inhibits Cu(I) binding (Fig. S3). However, in contrast to the internal motif, there are no discrete MDM clusters in this sequence; instead, seven methionine residues are roughly equally distributed across the length of the peptide, separated by residues that are mostly charged and polar. The N-terminal peptide undergoes a more profound structural rearrangement on Cu(I) binding and trimerization than does the internal motif (Fig. 6). We speculate that a specific knot-like structure of three peptides forms around a single Cu(I) ion, though further investigation will be required to understand the mechanism by which this occurs. However, untangling the question of the precise metal-binding mechanism of the N-terminal peptide is of particular import for understanding reflectin self-assembly, given that it is the N-terminal motif that best distinguishes the different reflectin isoforms expressed in a given tissue from one another. It is worth noting that a DLS experiment shows formation of larger aggregates of N-terminal peptides with increasing Cu(I) concentration, however these events are silent in ITC (Figs. S9, S8), suggesting that these aggregates occur via non-specific hydrophobic interactions of the trimers observed in ITC.

When we analyzed the phase separation of the internal motif as a function of peptide concentration, we observed that the fraction of peptide mass that separates into the dense phase behaves with bounded asymptotic growth as a function of peptide concentration. It was not technically possible to extend this experiment to higher peptide concentrations given the solubility of the peptide. However, in a living cell able to continuously add new protein to an equilibrating system, fitting this relation predicts that full phase separation into a volume-spanning material will occur at packing fractions ≥2%.

Therefore, we conclude that in coordination with Cu(I), the N-terminal motif forms specific trimers, while the internal motif undergoes a continuous self-assembly process that likely results in a network of branching chains. In the native system, a full-length protein consists of one N-terminal motif and 4–5 repeats of the internal motif. Therefore, the Cu(I)-binding properties of a full-length protein may allow for the self-assembly of a network formed of chains of internal motifs bound in antiparallel orientation with nodes in the network formed by trimers at the N-terminal ends of each protein (Fig. 10). These observations inform hypotheses for future work in elucidating how the squid reflectin system controls density, density fluctuation, phase separation, and geometry of the optically resonant arrays in the living system.

**Fig. 10:**
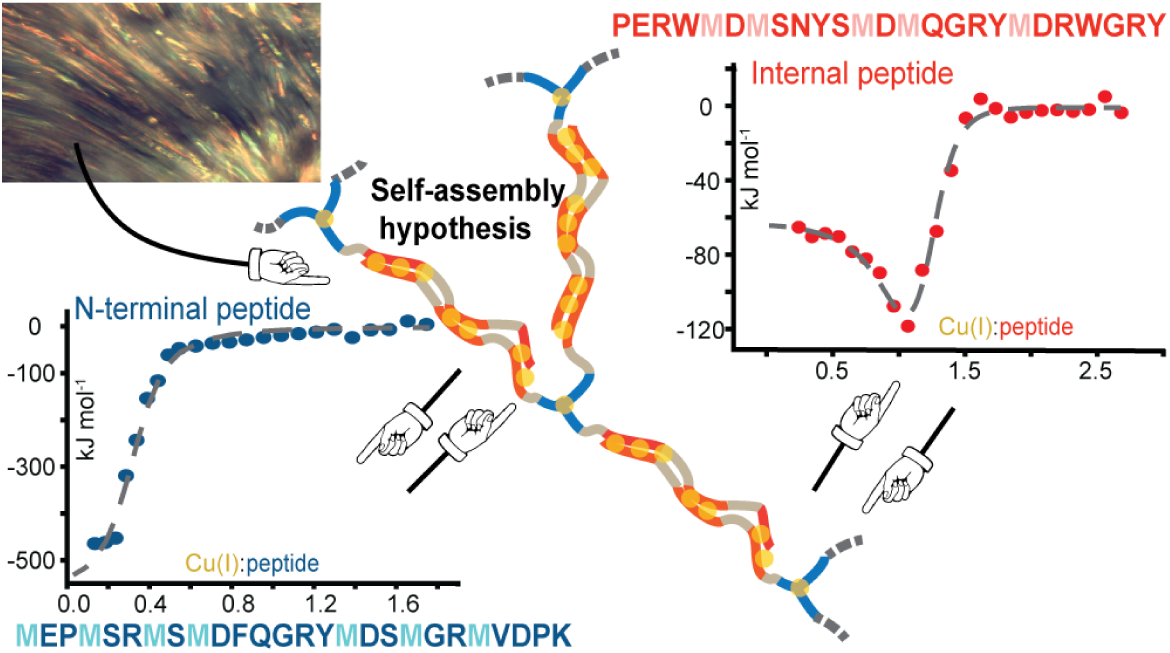
Structural self-assembly hypothesis of full-length reflectin proteins into a volume-spanning gel. In the assembly schematic, red regions represent internal motifs, blue regions represent N-terminal motifs, grey regions are aromatic-rich linkers, and yellow circles are Cu(I) ions.

This work also represents the discovery of two novel copper(I)-binding centers responsible for governing the self-assembly of a material. The squid reflectin system achieves self-assembling photonics in part via self-assembly through directed, specific copper cross-linking.

## Acknowledgement

The authors thank Dr. Fabian Menges of the Yale Chemical and Biophysical Instrumentation Center for technical assistance with MALDI-MS; Dr. Dan Asael of the Yale Metal Geochemistry and Geochronology Center for technical assistance with ICP-MS; Mengwen Shi and Charles Lomba for assistance with data analysis; Prof. Seth Herzon for assistance with NMR; and Profs. Enrique De La Cruz and Andrew Miranker, whose advice and suggestions improved our work.

## Supporting Information Available

- Figure S1: Comparison of yeast Mets motifs with reflectin motif.
- Figure S2: ^1^*H* NMR spectra of free acetonitrile in peptide with and without Cu(I).
- Figure S3: MALDI-MS data showing peptide-Cu(I) binding as a function of methionine oxidation.
- Figure S4: MALDI-MS data showing peptides bind more Cu(I) as a function of [Cu(I)].
- Figure S5: Ascorbic acid assay of motif and N-terminus peptides reacting with Cu(I).
- Figure S6: Bathocuproine disulfonic acid analysis of Cu in native tissue.
- Figure S7: CD control spectra of peptide in the presence of CuSO_4_ and NaCl.
- Figure S8: Evolving reaction kinetics of peptides derived from ITC with serial injections of Cu(I).
- Figure S9: DLS of peptides in a concentration series with Cu(I).
- Figure S10: ITC control spectrum of peptide in presence of FeSO_4_.
- Figure S11: Fractional binding events in MALDI-MS spectra of motif peptide.

