## Supplements for "The Cephalopod Reflectin Domain Specifically Chelates Cu(I), Inducing Photonic Assembly"

| | |
|---|---|
| yMets1 | MEGMNM |
| yMets2 | MNMDAM |
| yMets3 | MASMSMDAM |
| yMets4 | MSSMSMEAM |
| yMets5 | MSSMASMSM |
| yMets6 | MSGMSMSM |
| yMets7 | MSGMSGM |
| yMets8 | MDMDMSMGM |
| Refl | MDMSNYSMDM |

**Fig. S1: Comparison of yeast Mets motifs with reflectin motif** (Mets motifs from Rubino and colleagues[1])

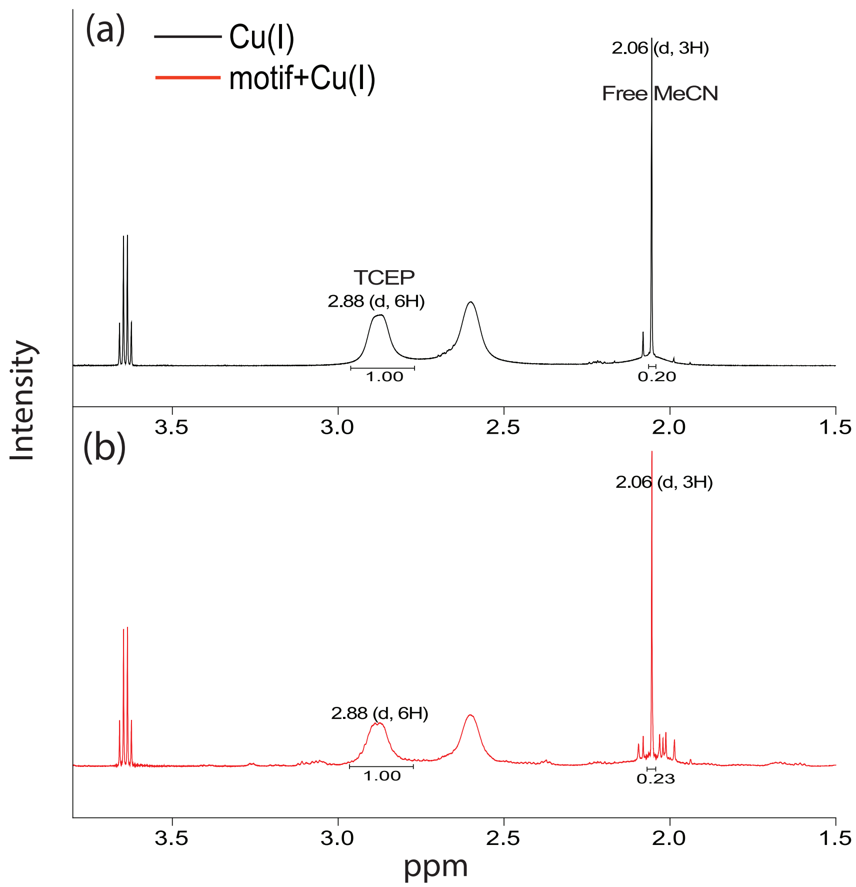

**Fig. S2:** $^1H$ NMR spectra of (a) 200$\mu$M $[Cu(CH_3CN)_4]PF_6$ in $D_2O$ with 2 mM TCEP (black) and (b) 100$\mu$M motif with 200$\mu$M $[Cu(CH_3CN)_4]PF_6$ in $D_2O$ with 2 mM TCEP(red). The free acetonitrile quantified with respect to the TCEP peak indicates all acetonitrile in the solution is free and unbound.

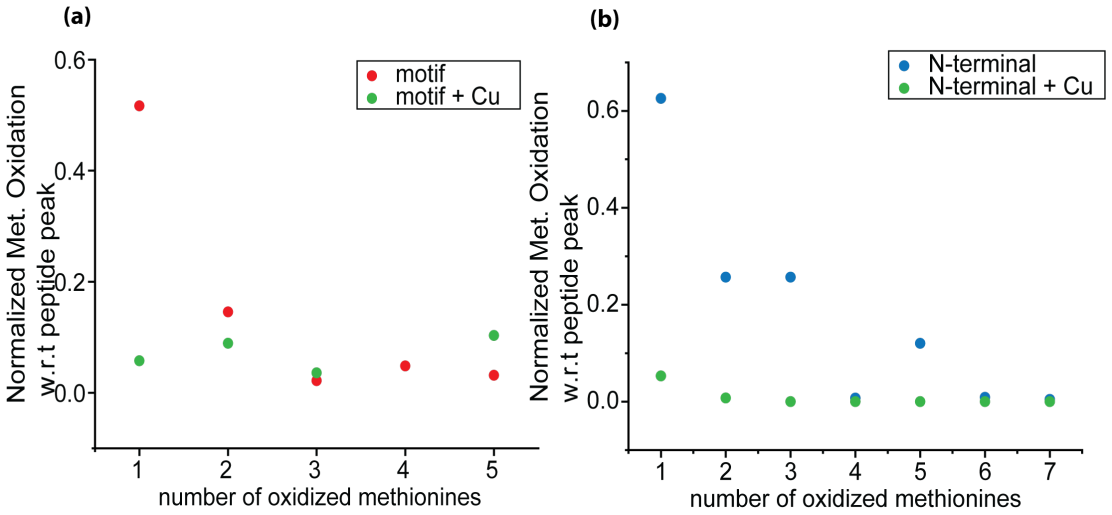

**Fig. S3: Peptide-Cu(I) binding as a function of methionine oxidation.** (a) Relative MALDI-MS peak intensities of the motif peptide when Cu(I)-bound (green points) and unbound peptide (red points) as a function of the number of oxidized methionine residues in the peptide. (b) Relative MALDI-MS peak intensities of the N-terminal peptide when Cu(I)-bound (green points) and unbound (blue points).

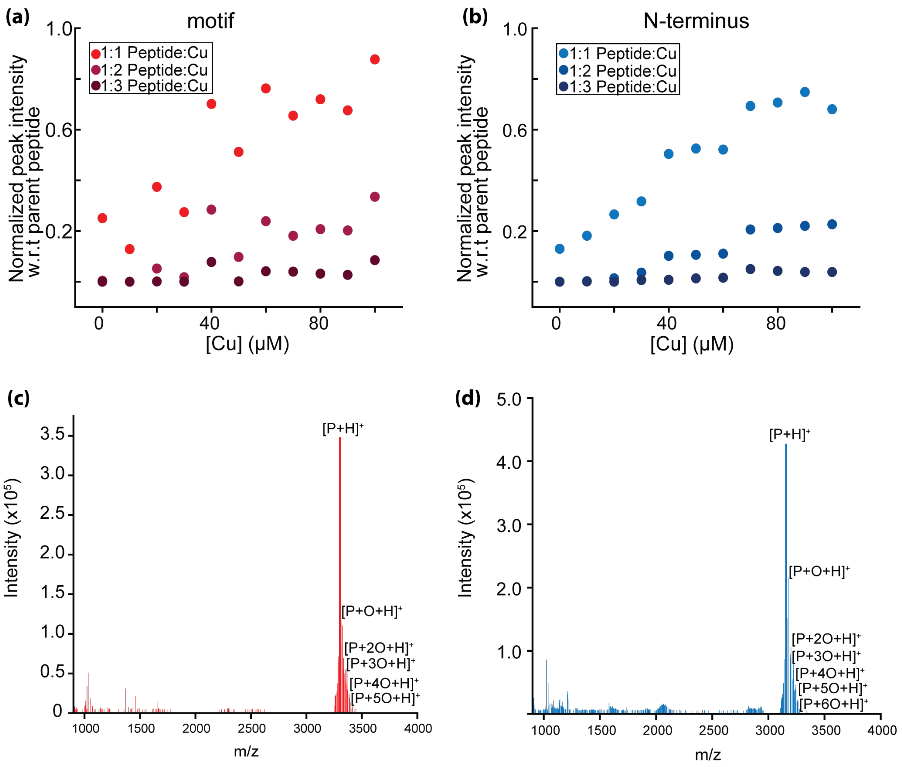

**Fig. S4: Peptides bind more Cu(I) as a function of [Cu(I)].** **(a)** Relative bound vs. unbound MALDI peak intensities of motif peptide in the presence of 1:1, 1:2, and 1:3 peptide:Cu. **(b)** Relative bound vs. unbound MALDI peak intensities of N-terminus peptide in the presence of 1:1, 1:2, and 1:3 peptide:Cu. **(c)** MALDI spectrum of motif peptide with Cu(II) and no ascorbic acid **(d)** MALDI spectrum of N-terminus peptide with Cu(II) and no ascorbic acid.

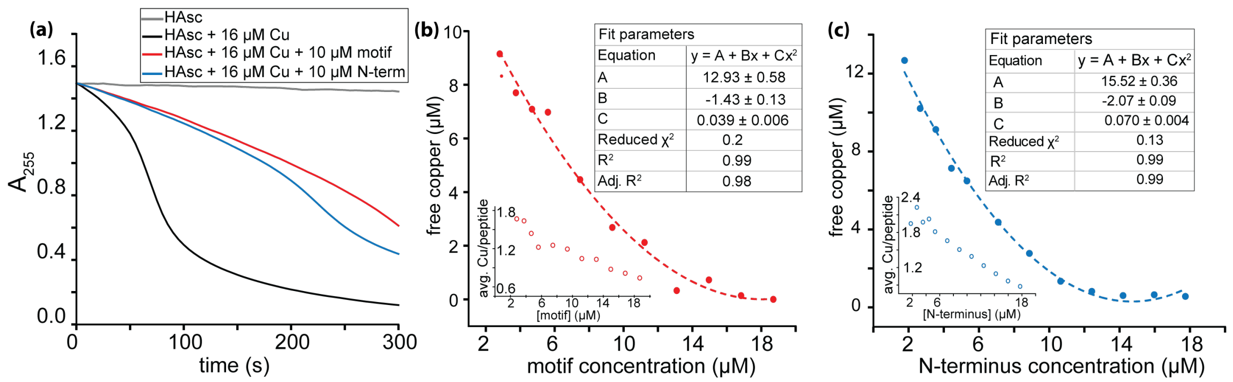

**Fig. S5: Ascorbic acid assay of motif and N-terminus peptides reacting with Cu(I) (a)** Absorbance at 255 nm of peptide-copper solutions and controls with time **(b)** Fitting of free copper as a function of the motif peptide concentration derived from data in **a** and the number of Cu bound per peptide as a function of peptide concentration (inset). **(c)** Fitting of free copper as a function of the N-terminus peptide concentration derived from data in **a** and the number of Cu bound per peptide as a function of peptide concentration (inset).

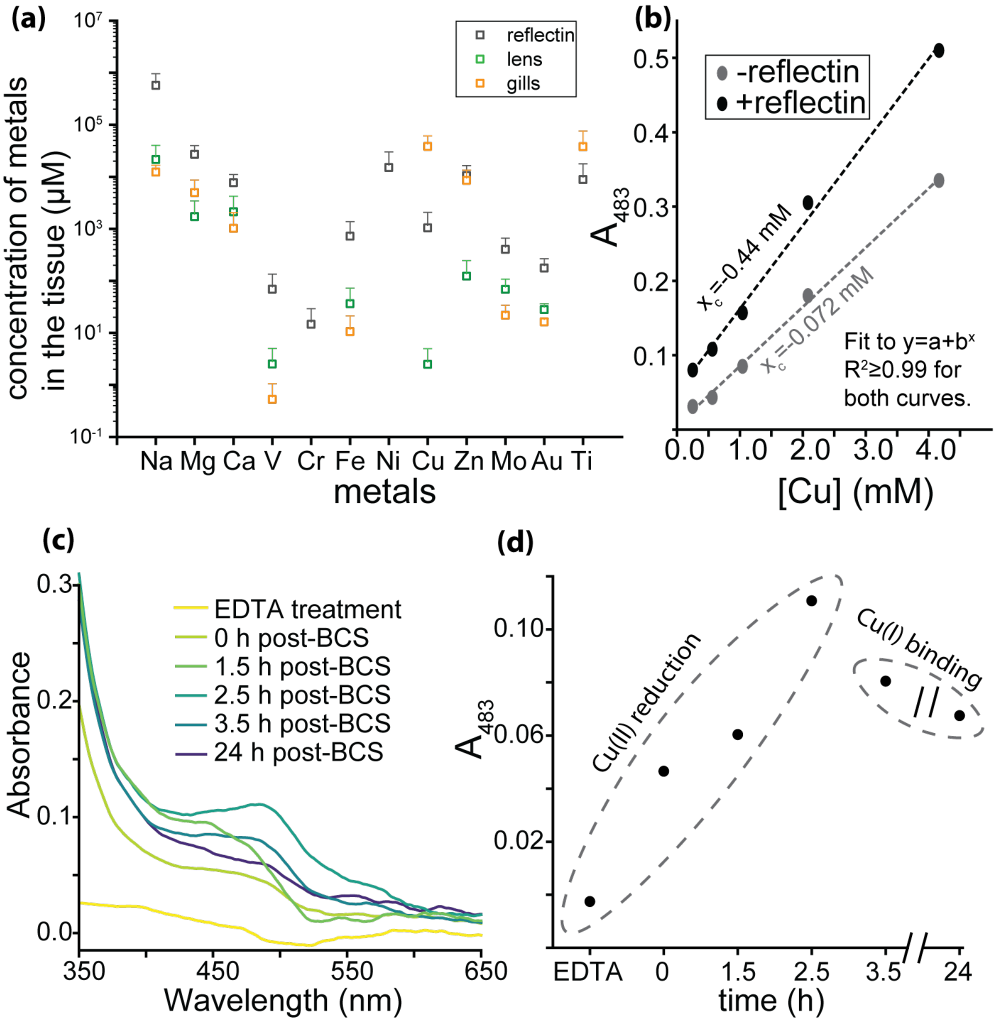

**Fig. S6: Analysis of Cu in native tissue.** **(a)** Transition-metal ICPMS of silver eye tissue, lens, and gill **(b)** A483 from Cu-BCS assay; A483 is systematically reduced in the presence of native protein and a linear fit indicates 8.9 Cu:1 reflectin protein. **(c)** BCS-A483 spectra of native reflectin tissue as a function of time show that reflectin can reduce Cu(II) and bind resulting Cu(I) **(d)** BCS valence test of native reflectin tissue treated to remove native metal ions. In the initial stage of the reaction, A483 increases in the presence of metal-free full-length reflectin, suggesting the protein reduces Cu(II) to Cu(I). In the next stage of the reaction, A483 decreases, suggesting the native protein out-competes BCS for Cu(I) removing it from solution with slower kinetics than BCS-Cu binding.

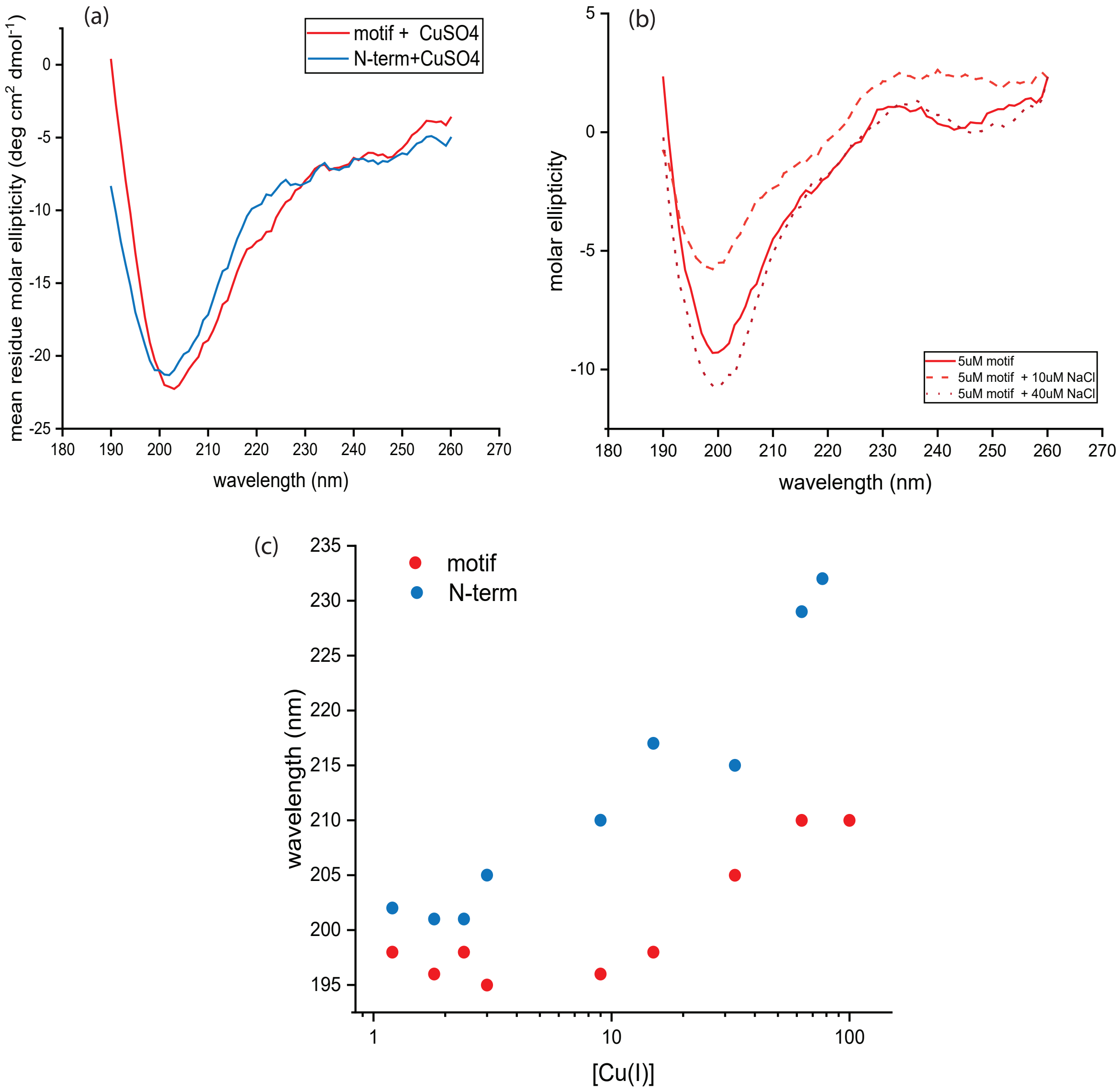

**Fig. S7:** (a) CD control spectra in the presence of $CuSO_4$, (b) CD spectra in the presence of NaCl. These controls confirm that CD structural changes observed in the presence of Cu(I) are not attributable to amyloid formation (a) or ionic strength (b). (c) The overall structural changes observed in the peptides during gradual Cu(I) titration.

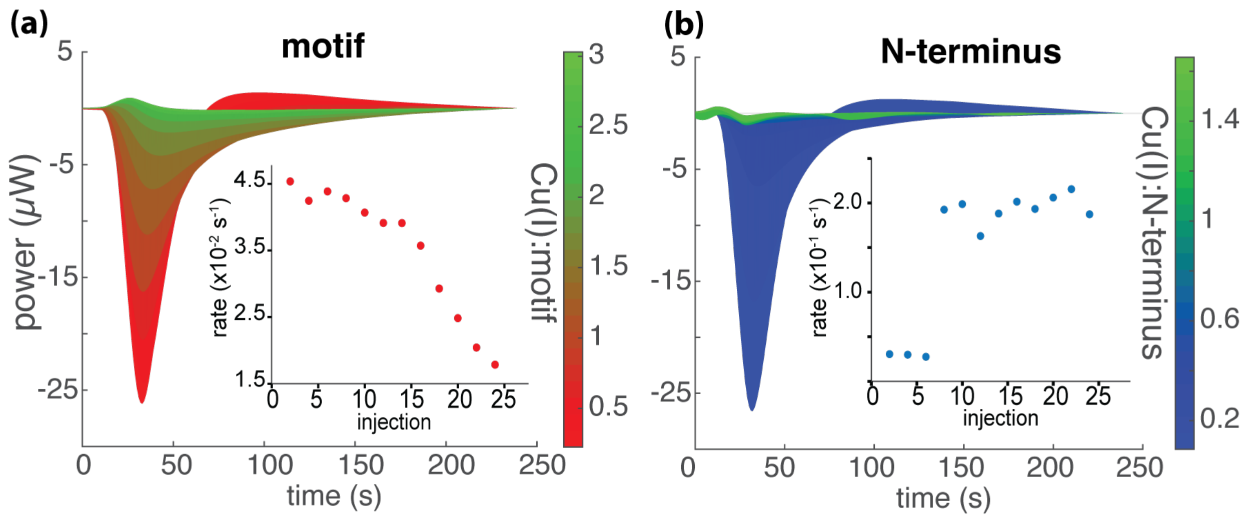

**Fig. S8: Evolving reaction kinetics derived from ITC with serial injections of Cu(I) reveal differently evolving reaction kinetics in the motif vs. the N-terminal peptide.** **(a)** Overlay of thermograms of Cu(I) titrated into motif peptide solution. At low motif peptide concentrations, Cu(I) binding is two times faster than at high motif peptide concentrations, where a slower event dominates the thermodynamic process. Inset: Reaction rate as a function of injection number. **(b)** Overlay of thermograms of Cu(I) titrated into N-terminal peptide Inset: Reaction rate as a function of injection number.

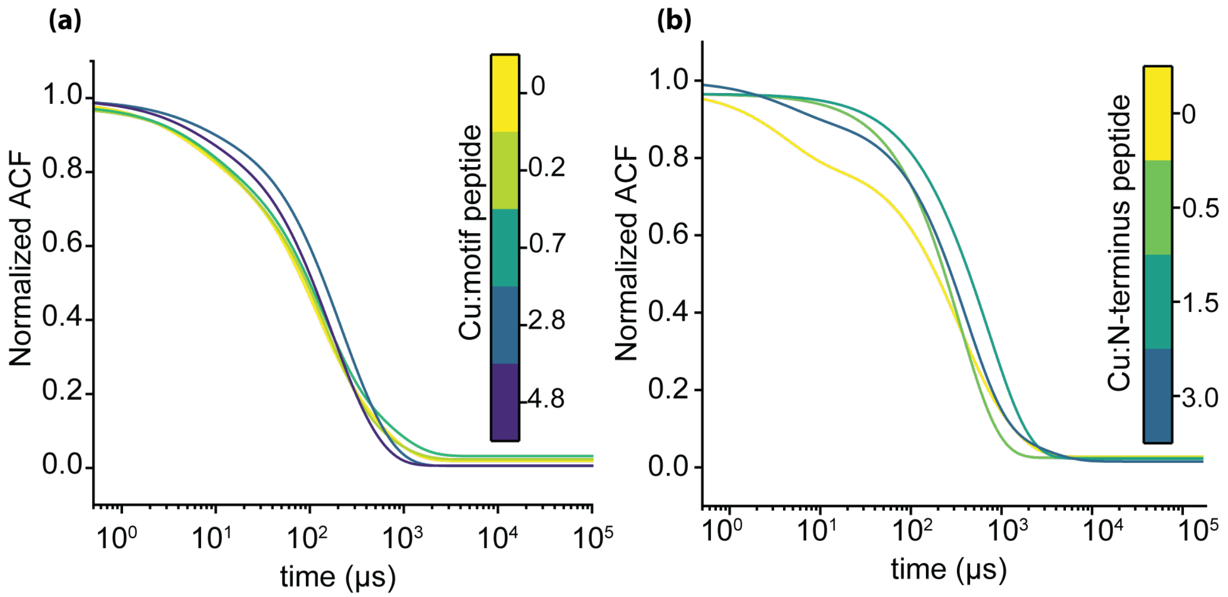

**Fig. S9: DLS of motif and N-terminus peptides in a concentration series with Cu(I) (a)** Time autocorrelation of dynamic light scattering of motif peptide in a range of molar ratios with Cu(I) **(b)** Time autocorrelation of dynamic light scattering of N-terminus peptide in a range of molar ratios with Cu(I)

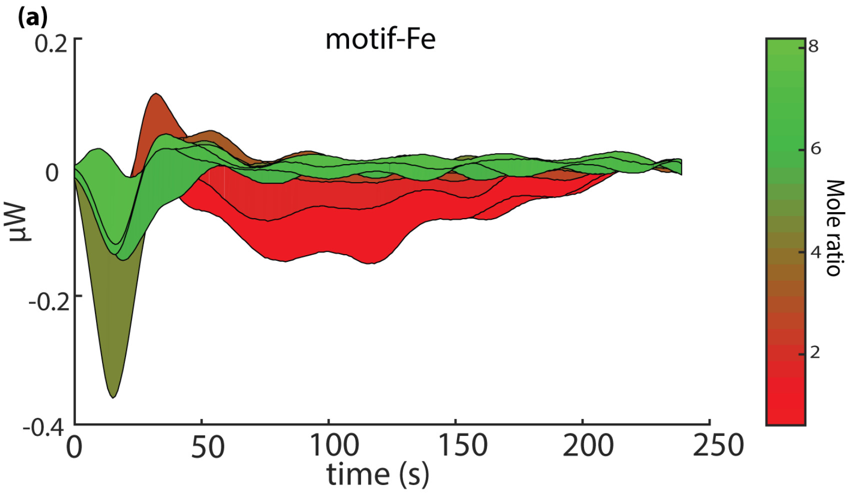

**Fig. S10:** (a) Overlay of titration curves generated for the motif in the presence of 1.5 mM $Fe^{2+}$ during a single experiment. The minimal raw heat signal, comparable to the noise level observed during the titration, indicates that the motif does not interact with $Fe^{2+}$

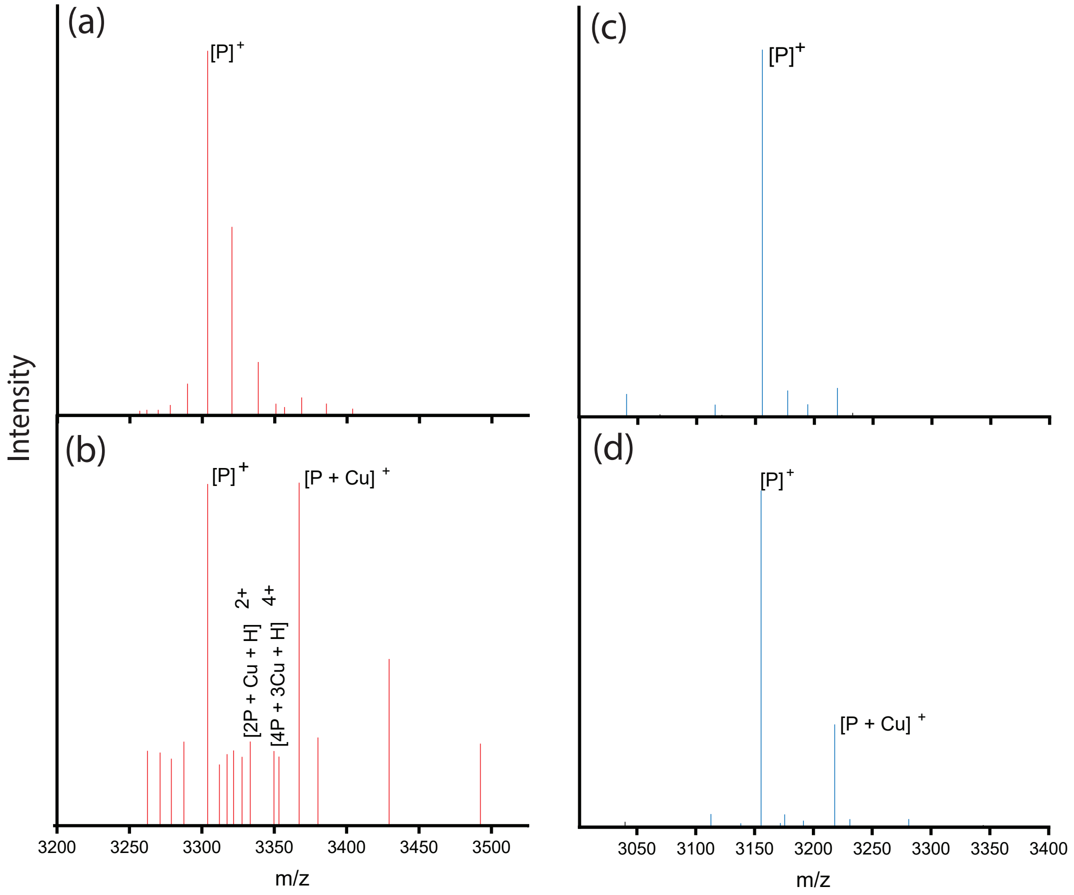

**Fig. S11:** Evidence of fractional binding events in MALDI-MS spectra of motif peptide (a) MALDI-MS spectrum of motif peptide in the absence of Cu(I). (b) MALDI-MS spectrum of N-terminal peptide in the absence of Cu(I) (c) MALDI-MS spectrum of motif peptide in the presence of Cu(I) (d) MALDI-MS spectrum of N-terminal peptide in the presence of Cu(I). Two or four peptides coordinated to Cu(I) at m/z of 3335.7 $[2P + Cu + H]^{2+}$ and 3351.2 $[4P + 3Cu + H]^{4+}$ as marked on the figure
